# Cortico-autonomic local arousals and heightened somatosensory arousability during NREM sleep of mice in neuropathic pain

**DOI:** 10.1101/2021.01.04.425347

**Authors:** Romain Cardis, Sandro Lecci, Laura M.J. Fernandez, Alejandro Osorio-Forero, Paul Chu Sin Chung, Stephany Fulda, Isabelle Decosterd, Anita Lüthi

**Affiliations:** Department of Fundamental Neurosciences, Rue du Bugnon 9, CH-1005 Lausanne; Pain Center, Department of Anesthesiology, Lausanne University Hospital (CHUV), Lausanne, Switzerland; Sleep and Epilepsy Center, Neurocenter of Southern Switzerland, Civic Hospital (EOC) of Lugano, Lugano, Switzerland

## Abstract

Chronic pain patients frequently suffer from sleep disturbances. Improvement of sleep quality alleviates pain, but neurophysiological mechanisms underlying sleep disturbances require clarification to advance therapeutic strategies. Chronic pain causes high-frequency electrical activity in pain-processing cortical areas that could disrupt the normal process of low-frequency sleep rhythm generation. We found that the spared-nerve-injury (SNI) mouse model, mimicking human neuropathic pain, had preserved sleep-wake behavior. However, when we probed spontaneous arousability based on infraslow continuity-fragility dynamics of non-rapid-eye-movement sleep (NREMS), we found more numerous local cortical arousals accompanied by heart rate increases in hindlimb primary somatosensory, but not in prelimbic, cortices of SNI mice. Closed-loop mechanovibrational stimulation revealed higher sensory arousability in SNI. Sleep in chronic pain thus looked preserved in conventional measures but showed elevated spontaneous and evoked arousability. Our findings develop a novel moment-to-moment probing of NREMS fragility and propose that chronic pain-induced sleep complaints arise from perturbed arousability.

## Introduction

Pain causes functional impairment, displeasure and stress and can impede even the simplest daily life routines, including sleep. If not treated, pain has the ability to outlast its original cause, producing chronic pain that is generally difficult to treat (Finnerup *et al*., 2015; Treede *et al*., 2019). Current estimates are that more than two out of three individuals suffering from chronic pain also show diverse symptoms characteristic for insomnia disorders, such as lower sleep efficiency, more time awake after sleep onset and frequent brief awakenings during the night (Bjurstrom & Irwin, 2016; Mathias *et al*., 2018). The relation between chronic pain and sleep disruptions is complex and bidirectional, but accurate assessment of sleep problems is considered critical to antagonize the perpetuation of pain (Bjurstrom & Irwin, 2016). Therefore, key mechanisms associating pain with sleep disturbance need to be clarified.

Animal models of chronic pain that mimic clinical symptoms of human patients have been critical to understand the pathophysiological mechanisms producing chronic pain states (Burma *et al*., 2017). Neuropathic pain is caused by damage to the somatosensory nervous system (Finnerup *et al*., 2021) and induced in rodents by surgically lesioning peripheral nerves, such as the sciatic nerve (Decosterd & Woolf, 2000; Bourquin *et al*., 2006). Neuropathic pain causes maladaptive structural and functional remodeling of the central and peripheral nervous systems, shifting brain circuits towards pain hypersensitivity and aversive behavioral states (Kuner & Kuner, 2020). Hyperexcitability and an abnormal activity in a broad range of gamma frequencies (30—100 Hz) in pain-processing cortical areas were found to be primary culprits for the elevated sensitivity to painful stimuli and for aversive behaviors (Tan *et al*., 2019), a finding that is in line with observations in human (Ploner *et al*., 2017). In contrast to these advances, sleep studies on chronic pain models are scarce, used relatively simple sleep measures, and produced variable results (Andersen & Tufik, 2003; Kontinen *et al*., 2003; Tokunaga *et al*., 2007; Cardoso-Cruz *et al*., 2011; Leys *et al*., 2013). Therefore, it is currently open whether these animal models are also suited to address the sleep complaints of human patients.

One possibility is that current approaches have so far failed to uncover the full profile of the sleep disruptions caused by chronic pain. Studies in insomnia disorders indeed suggest that changes in traditional sleep parameters often seem not in line with the severity of the sleep complaints (Feige *et al*., 2013; van Someren, 2020). Standard polysomnography describes sleep as a sequence of discrete states and distinguishes between non-rapid-eye-movement sleep (NREMS) and rapid-eye-movement sleep (REMS), with the former further subdivided into transitional (N1), light (N2) and deep (N3) stages (Iber *et al*., 2007). Many reports on human patients find little change in the absolute or relative times spent in these stages and/or their principal spectral characteristics (Salin-Pascual *et al*., 1992; Perlis *et al*., 2001b; Buysse *et al*., 2008; Wei *et al*., 2017; Feige *et al*., 2018; Christensen *et al*., 2019; Lecci *et al*., 2020). Instead, cortical activity patterns are abnormally enriched in the alpha (8—12 Hz) (Krystal *et al*., 2002; Riedner *et al*., 2016), beta (18—30 Hz) (Krystal et al., 2002; Spiegelhalder *et al*., 2012; Maes *et al*., 2014; Riedner et al., 2016; Lecci et al., 2020) and/or low-gamma bands (30—45 Hz) (Perlis et al., 2001b; Lecci et al., 2020), in one or more NREMS stages and/or in REMS (Spiegelhalder et al., 2012; Christensen et al., 2019; Lecci et al., 2020) and/or in restricted brain areas (St-Jean *et al*., 2012; Riedner et al., 2016; Lecci et al., 2020). Such high-frequency electrical rhythms during sleep are part of a physiological state referred to as “hyperarousal” (Feige et al., 2013; van Someren, 2020; Vargas *et al*., 2020) that has been related to less restorative sleep (Moldofsky *et al*., 1975; Krystal & Edinger, 2008), to misperceiving sleep as wakefulness (Perlis et al., 2001b; Lecci et al., 2020), and to higher heart rates (Maes et al., 2014), all of which are key features of insomnia disorders in humans. Other studies applied various metrics and proposed more spontaneous arousals and/or easier wake-ups in response to sensory stimulation (Parrino *et al*., 2009; Forget *et al*., 2011; Wei et al., 2017; Feige et al., 2018). Taken together, the presence of high-frequency electrical activity, combined with diverse measures of arousability, has been useful in clarifying the pathophysiological mechanisms underlying insomnia disorders. To date, however, such measures have not been applied to study sleep in chronic pain in humans and mice, and there is still a paucity of comparative studies between sleep in chronic pain patients and in primary insomniacs (Bjurstrom & Irwin, 2016). Therefore, it is currently unclear whether chronic pain is accompanied by high-frequency cortical activity during NREMS and whether this affects spontaneous and evoked arousability (Mathias et al., 2018; Kuner & Kuner, 2020).

This study pursues this question through implementing a real-time tracking method for spontaneous and evoked arousability in the mouse spared-nerve-injury (SNI) model of neuropathic pain (Bourquin et al., 2006). We start from previously described fragility-continuity dynamics of NREMS in mice and humans that indicate variable arousability on the ∼50-sec time scale (Lecci *et al*., 2017). The fragility-continuity dynamics are present while NREMS remains polysomnographically continuous and manifest in fluctuating activity of several brain and peripheral parameters, notably in the power of sleep spindles (10—15 Hz) in the global EEG and the local field potential (LFP) signals. On this close-to-minute timescale, we show here that we are capable of tracking spontaneous and evoked arousability across NREMS in the resting (light) phase. We find that the sleep disruptions in SNI animals concern both, altered spontaneous and evoked arousability. In particular, we identify a novel, previously undescribed cortico-autonomic arousal that pairs EEG desynchronization with increased heart rate and that occurs more frequently in SNI animals.

## Results

### SNI mice show normal sleep-wake behavior

SNI mice and Sham controls (n = 18 each) were first analyzed for their sleep-wake behavior using standard polysomnography. EEG/EMG measurements were carried out prior to SNI and Sham surgery and at post-surgical days 22-23 (D20+), a time point at which chronic pain is established (Decosterd & Woolf, 2000; Bourquin et al., 2006). At both time points, recordings lasted for 48 h under undisturbed conditions. Based on these data, SNI and Sham controls spent similar amounts of time asleep in the 12 h-light and dark phases, during both baseline and at D20+ (***Figure 1A***). Both treatment groups showed minor increases (2.4-3 %) in NREMS time at the expense of wakefulness at D20+ compared to baseline in both dark and light phases (mixed-model ANOVAs with factors ‘treatment’ and ‘day’, p = 0.0013 in light phase, p = 8.5×10^−6^ in dark phase for ‘day’, p > 0.8 for ‘treatment’ for either light or dark phase, no interaction). Moreover, cumulative distributions of NREMS and REMS bout lengths at D20+ were similar for Sham and SNI, with only a minor shift toward smaller values in SNI for both NREMS (***Figure 1B***, −2.3 s; Kolmogorov-Smirnov (KS) test, p = 0.015) and REMS (−3 s; KS test, p = 0.026). When subdivided into short, intermediate and long bouts for the light phase, there were no significant differences between Sham and SNI for both NREMS and REMS (***Figure 1B***, mixed-model ANOVAs for ‘treatment’ x ‘bout length’, p = 0.79 for NREMS, p = 0.23 for REMS).

**Figure 1.**
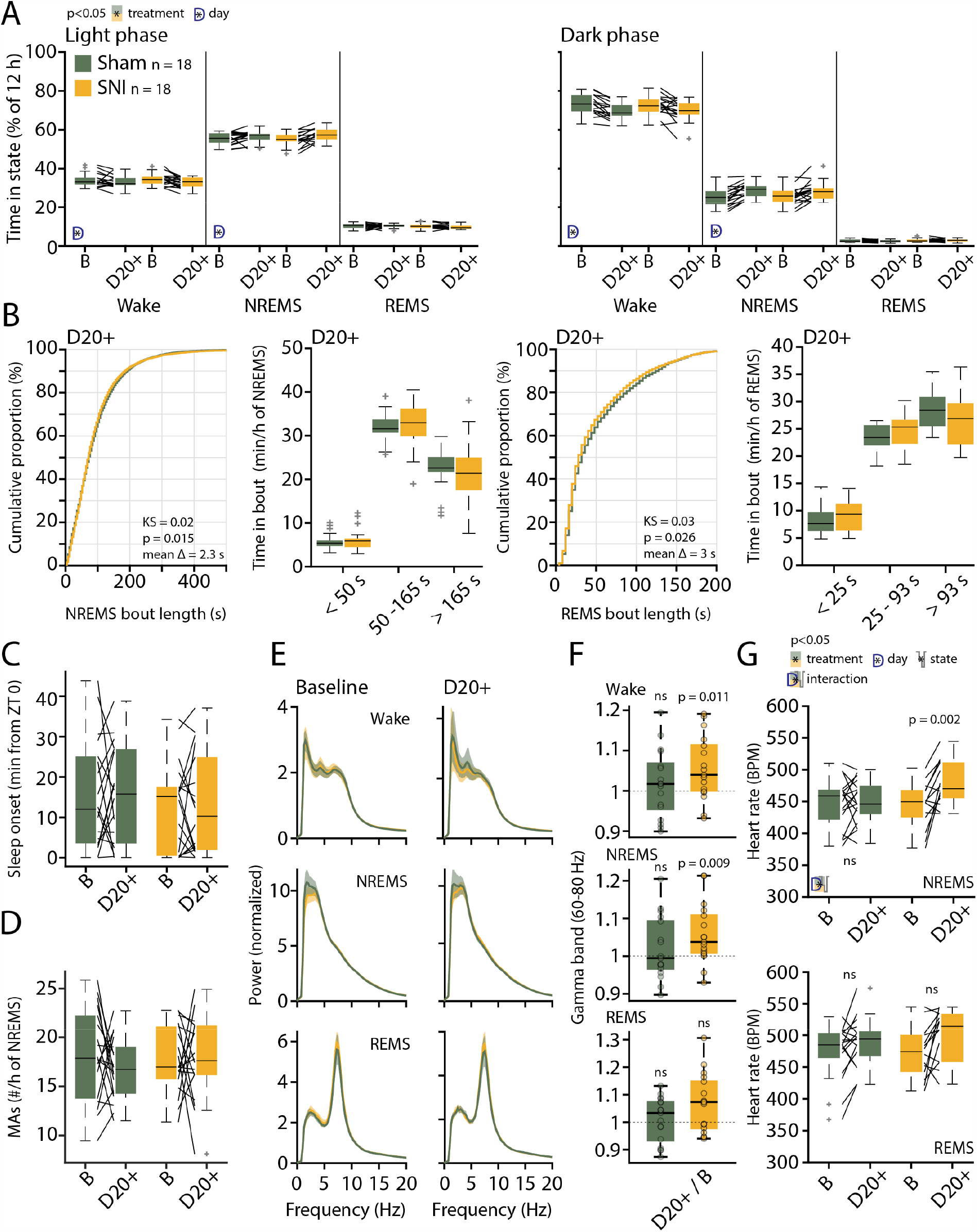
Preserved sleep-wake behavior and spectral properties in SNI animals. (**A**) Mean percentage of total time spent in the three main vigilance states for Sham (n = 18) and SNI (n = 18) animals in light (left) and dark phase (right) in baseline (B) and at day 20+ (D20+) after surgery. Black lines connect data from single animals. The significant main effects and interactions from the ANOVAs are shown using pictograms for factors and interaction. *Post-hoc* tests were done once interactions were significant. Mixed-model ANOVA for non-rapid-eye-movement sleep (NREMS) in light phase: F_(1,34)_ = 0.048, p = 0.82 for ‘treatment’; F_(1,34)_ = 12.24, p = 0.0013 for ‘day’; F_(1,34)_ = 2.5, p = 0.12 for interaction. Mixed-model ANOVA for NREMS in dark phase: F_(1,34)_ = 0.009, p = 0.92 for ‘treatment’; F_(1,34)_ = 27.409, p = 8.5×10^−6^ for ‘day’; F_(1,34)_ = 0.379, p = 0.54 for interaction. There were no significant effects in rapid-eye-movement sleep (REMS). (**B**) Bout size cumulative distribution, (with Kolmogorov Smirnov (KS) test results) and time spent in short, intermediate and long bouts for NREMS (left) and REMS (right) between Sham and SNI at D20+. Mixed-model ANOVA for time in bouts (log transform for normality criteria) for NREMS: F_(2,68)_ = 0.23, p = 0.79 for interaction, for REMS: F_(2,68)_ = 1.5, p = 0.23 for interaction. (**C**) Mean latency to sleep onset (first consolidated NREMS). Mixed-model ANOVA F_(1,34)_ = 0.7, p = 0.4 for ‘treatment’; F_(1,34)_ = 0.006, p = 0.94 for ‘day’; F_(1,34)_ = 0.021, p = 0.88 for interaction. (**D**) Number of microarousals (MAs) per h of NREMS in light phase; Mixed-model ANOVA F_(1,34)_ = 0.067, p = 0.8 for ‘treatment’; F_(1,34)_ = 0.63, p = 0.43 for ‘day’; F_(1,34)_ = 0.61, p = 0.44 for interaction. (**E**) Normalized power spectrum for Sham and SNI for wake, NREMS and REMS at baseline (left) and D20+ (right). Shaded errors are 95 % confidence intervals (CIs) of the means. (**F**) High-gamma power (60—80 Hz) for D20+ relative to baseline (from the spectra shown in E). One-sample *t*-test for wake in Sham: t_(16)_ = 0.86, p = 0.4; for wake in SNI: t_(16)_ = 2.8, p = 0.011. One-sample *t*-test for NREMS in Sham: t_(16)_ = 0.97, p = 0.34; for NREMS in SNI: t_(16)_ = 2.96, p = 0.0092. One-sample *t*-test for REMS in Sham: t_(16)_ = 0.46, p = 0.64; for REMS in SNI: t_(16)_ = 2.62, p = 0.0184. α = 0.0167. (**G**) Heart rate in NREMS (top) and REMS (bottom) from animals with suitable EMG signal, Sham (n = 17) and SNI (n = 14). Mixed-model three-way ANOVA: F_(1,29)_ = 0.5, p = 0.47 for ‘treatment’; F_(1,29)_ = 70.4, p = 3×10^−9^ for ‘state’, F_(1,29)_ = 9.79, p = 0.003 for ‘day’; F_(1,1,29)_ = 5.37, p = 0.02 for interaction between the three factors; *post-hoc* paired *t*-test in NREMS for baseline *vs* D20+ in Sham: t_(16)_ = −0.4, p = 0.69; SNI: t_(13)_ = − 3.75, p = 0.002; paired *t*-test in REMS for baseline *vs* D20+ in Sham: t_(16)_ = −1.9, p = 0.07; SNI: t_(13)_ = −2.4, p = 0.027; α = 0.0125.

Furthermore, sleep onset latency (***Figure 1C***) and NREMS fragmentation by brief movement-associated microarousals (MAs, defined in mouse as <= 16 s awakenings accompanied by movement activity seen in the EMG, measured over 48 h) (***Figure 1D***) (Franken *et al*., 1999), were not altered by treatment or time post-surgery (mixed-model ANOVA with factors ‘treatment’ and ‘day’, for sleep onset latency, p = 0.42 and p = 0.94, no interactions; for number of MAs, p = 0.79 and p = 0.43, no interactions).

We next investigated the mean spectral properties of each vigilance state through constructing normalized power spectral densities (Vassalli & Franken, 2017) for the full 48 h-long recordings. Both NREMS and REMS showed the respective characteristic spectral peaks at delta (1—4 Hz) and at theta frequencies (5—10 Hz), respectively. These were indistinguishable between the two groups of animals and from baseline to D20+ (***Figure 1E***).

We specifically evaluated power in the high-gamma frequency (60—80 Hz) range, a frequency band linked to pain sensations when optogenetically induced in mouse (Tan et al., 2019). We found that relative gamma power was increased in SNI at D20+ compared to baseline both in wake and in NREMS (***Figure 1F***, 1-sample *t*-test for wake and NREMS in SNI, p = 0.011 and p = 0.0092, respectively). The heart rate was also higher in NREMS of SNI animals at D20+ compared to baseline (***Figure 1G***, mixed-model ANOVA with factors ‘treatment’ x ‘state’ x ‘day’ with interaction, p = 0.02, *post-hoc* paired *t*-test for SNI in NREMS, p = 0.002, with Bonferroni-corrected α = 0.0125). A tendency was also evident in REMS, during which heart rate was already elevated (***Figure 1G***, effect of ‘state’ in the ANOVA, p = 0.003, paired *t*-test in SNI in REMS, p = 0.027). There were no correlations between relative changes in gamma power and alterations in sleep architecture in individual mice (change in the number of MAs per h of NREMS x change in gamma power; pairwise linear correlation R^2^ = 0.09, p = 0.08; change in total NREMS time x change in gamma power; R^2^ = 0.02, p = 0.36).

These data indicate that SNI animals do not suffer from major alterations in sleep-wake behaviors. Still, pain-related pathological changes in brain and periphery continued to be present in sleep. This is consistent with a state of “hyperarousal” whereby high-frequency power components are disproportionately elevated during sleep that is normally dominated by low-frequency rhythms (van Someren, 2020; Vargas et al., 2020), and where heart rate also remains elevated. As such alterations could affect arousability, we asked when and where in the brain this abnormal activity appeared. Furthermore, we developed an approach to systematically quantify alterations in both, spontaneous and evoked, types of arousability from NREMS during the resting phase.

### The 0.02 Hz-fluctuation allows to probe variations in spontaneous arousability during NREMS

Arousability in sleeping rodent, measured via external stimuli or through spontaneous arousals, changes across the night (Neckelmann & Ursin, 1993; Wimmer *et al*., 2012), and with variations in sleep pressure (Franken et al., 1999). For NREMS in early phases of the resting phase, we described a 0.02 Hz-fluctuation during NREMS that provides a minute-by-minute time raster to measure arousability driven by sensory stimuli. This fluctuation subdivides NREMS bouts into ∼25 s-long periods of continuity and fragility that show low and high sensory-evoked arousability, respectively (Lecci et al., 2017; Yüzgeç *et al*., 2018). To evaluate the utility of this fluctuation for measures of spontaneous arousability across the entire light phase, we first tested whether MAs associated with muscular activity (***Figure 2A***), well-established correlates for spontaneous arousability, were phase-locked to the 0.02 Hz-fluctuation in healthy mice (n = 30 mice with 9,476 MAs). The onset of MAs coincided with declining or low sigma power levels that followed a pronounced sigma power peak (***Figure 2B, C***), which is characteristic for a fragility period (Lecci et al., 2017; Fernandez & Lüthi, 2020). A spectral band typical for NREMS, such as delta (1—4 Hz) power, showed a rapid decline preceding the MAs, indicating the momentary interruption of NREMS. The phase values of the 0.02 Hz-fluctuation, calculated via a Hilbert transform (***Figure 2 – figure supplement 1***), showed that MA onset times clustered around a mean preferred phase of 151.6° ± 1.1°, with 180° representing the sigma power trough (Rayleigh test, p < 1×10^−16^). The majority of MAs (89 %) was clustered between 90—270°, which narrows the fragility period to the low values of sigma power around the trough (Lecci et al., 2017). The phase-locking was also observed when time points at 4, 8 and 12 s before the onset of a MA were quantified (***Figure 2D***). This shows that the onset of the fragility period preceded the MA. Fragility periods thus constitute moments during which MAs preferentially occur.

**Figure 2.**
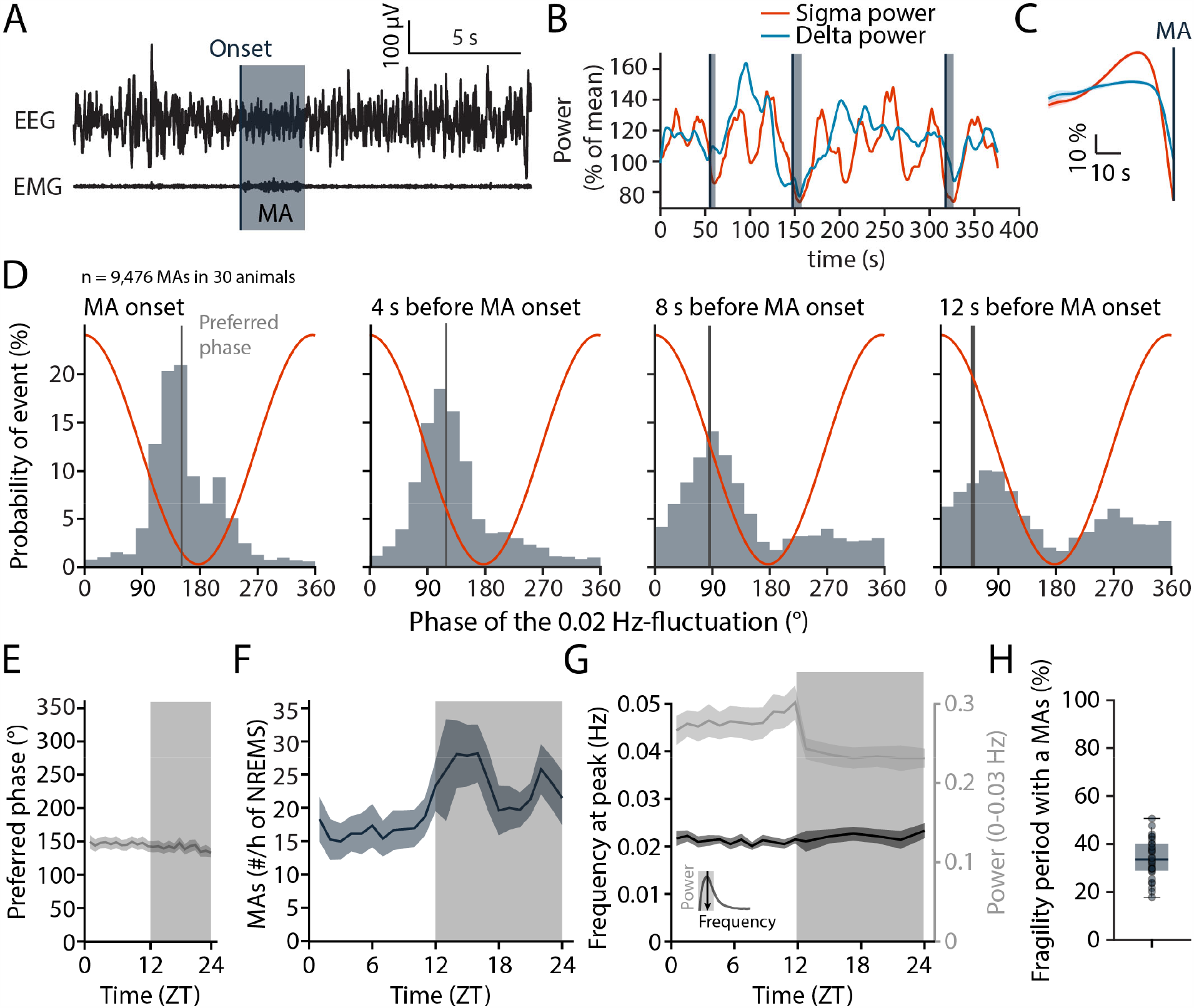
MAs, indicating spontaneous arousability, are time-locked to the trough of the 0.02 Hz-fluctuation, corresponding to the NREMS fragility period. (**A**) Example of a MA (underlain in grey) defined by both a desynchronization in the EEG (top) and a burst of EMG activity (bottom) of length <16 s. The MAs were scored visually, and the onset was set at the beginning of phasic EMG activity (black vertical line). (**B**) Continuous sigma (red; 10—15 Hz) and delta (blue; 1—4 Hz) power in a NREMS period with MAs (grey-shading as in panel A). Note their occurrence during descending / low values of sigma power. (**C**) Mean sigma and delta power dynamics preceding the onset of a MA. (**D**) Histograms of the phase angle values of the 0.02 Hz-fluctuation at specific time points relative to the onset of a MA. The red line represents the corresponding phase of the fluctuation at each bin. A total of n = 9,476 MAs from 30 un-operated C57Bl/6J animals were included across the light-dark cycles for all analyses in this figure. Rayleigh tests and preferred phases ± 95% CI for MA onset: z = 4214.7, p < 1×10^−16^, m = 151.5 ± 1.1°; for 4 s before MA onset: z = 3489, p < 1×10^−16^, m = 119.1 ± 1.2°; for 8 s before MA onset: z = 1416, p < 1×10^−16^, m = 85.3 ± 2.0°; for 12 s before MA onset: z = 548.163, p < 1×10^−16^, m = 50.1 ± 3.3°. (**E**) Preferred phase of the 0.02 Hz-fluctuation at MA onset across time-of-day in hourly bins (dark phase, shaded, ZT, Zeitgeber time). (**F**) Density of MAs (per h of NREMS) across the light-dark cycle. (**G**) Parameters of the 0.02 Hz-fluctuation (frequency at peak and power, see inset for illustration) across the light-dark cycle. (**H**) Proportion of fragility periods (corresponding to values from 90 to 270°, see panel D) containing a MA.

These phase relations persisted for all 1-h intervals across time-of-day (***Figure 2E***), although the density of MAs showed a characteristic increase towards the end of the light phase and was higher during the dark phase (***Figure 2F***). The peak frequency of the 0.02 Hz-fluctuation also remained relatively constant, with a minor decrease in power during the dark phase (***Figure 2G***). Across the 24-h cycle, a median of 33.6 % of all fragility periods were accompanied by a MA (***Figure 2H***). In sum, fragility periods are permissive windows for MAs. This means that MAs appeared predominantly during fragility periods, while a majority of fragility periods occurred with NREMS remaining consolidated.

### NREMS in SNI conditions shows normal phase-coupling of MAs to the 0.02 Hz-fluctuation

We next evaluated SNI and Sham animals regarding the MAs and their coupling to the 0.02 Hz-fluctuation. The 0.02 Hz-fluctuation was not different between Sham and SNI (n = 18 for both groups) across the light phase. Thus, neither its amplitude nor frequency (***Figure 3A-C***), or, equivalently, the number of its cycles per h of NREMS, were different between the groups (***Figure 3D***). The phase-coupling to MAs was also unaltered (***Figure 3E***, mean angle ± 95% CI: 152.3 ± 1.4 for Sham and 150.4 ± 1.3 for SNI) and the distribution of fragility periods containing transitions to MAs, to REMS, or with continuation into NREMS was indistinguishable (***Figure 3F***).

**Figure 3.**
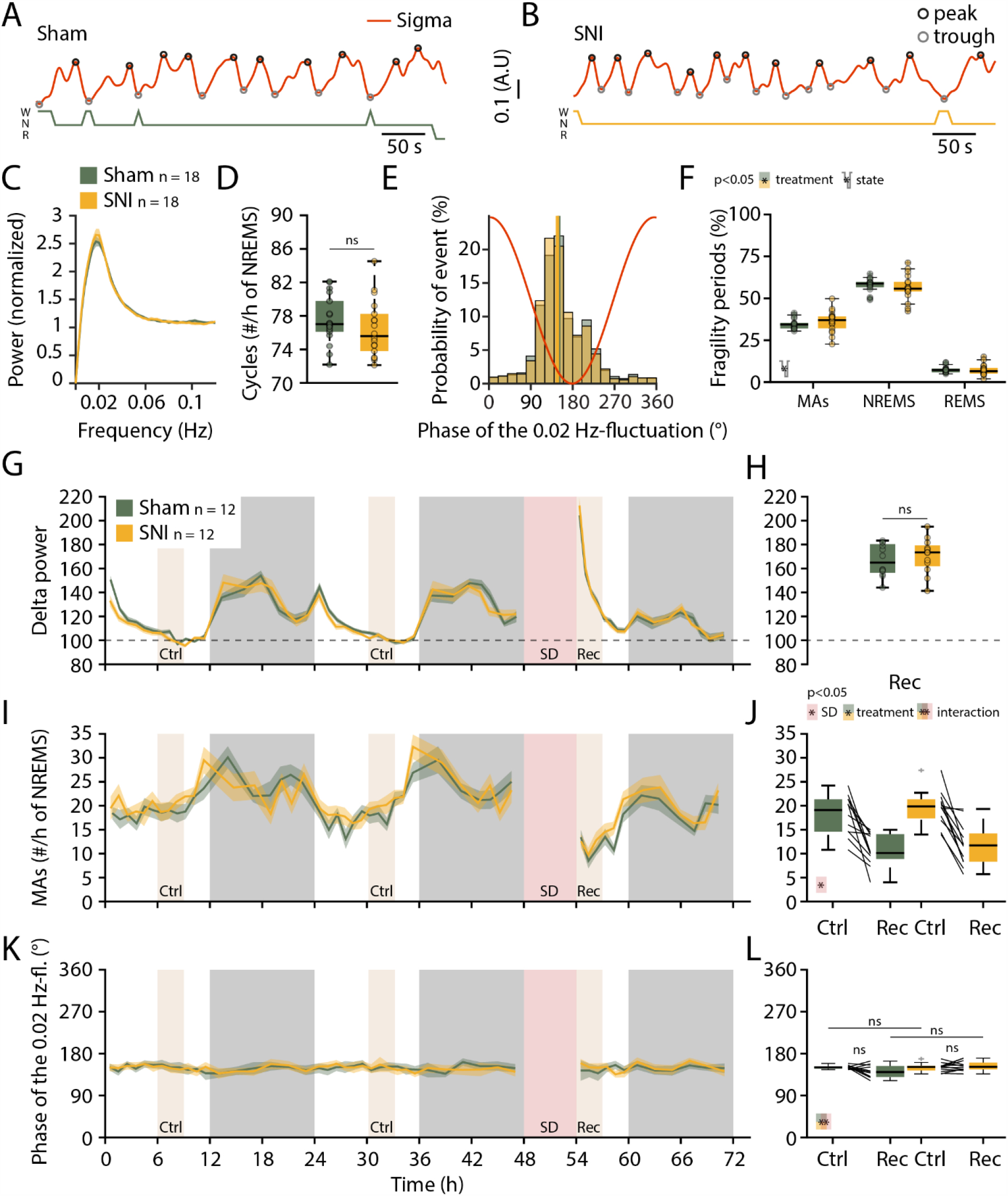
The 0.02 Hz-fluctuation and its relationship to MAs were preserved in SNI animals, even when sleep pressure was high. (**A-B**) Representative traces of sigma power dynamics in a Sham (A) and a SNI (B) animal. Hypnograms shown below for W, wakefulness; N, NREMS; R, REMS. The circles represent the individual cycle detection used in D and F (see methods). (**C**) Power in the infraslow range for Sham and SNI (n = 18 each); shaded areas represent 95% CIs. (**D**) Another measure of the 0.02 Hz-fluctuation, calculated as number of cycles per h of NREMS. Data are shown for D20+ only, but baseline data points were considered for statistical analysis. Mixed-model ANOVA: F_(1,34)_ = 0.012, p = 0.91 for ‘treatment’; F_(1,34)_ = 0.003, p = 0.95 for ‘day’; F_(1,34)_ = 2.17, p = 0.14 for the interaction. (**E**) Histograms of the phase values of the 0.02 Hz-fluctuation at MA onset for Sham and SNI, same as Figure 2D. Vertical lines denote mean direction ± 95% CI for Sham: 152.3 ± 1.4; SNI: 150.4 ± 1.3. (**F**) Proportion of fragility periods (defined by 0.02 Hz-fluctuation phase values of 90—270°) containing a MA, continuing into NREMS or containing a transition to REMS. Mixed-model ANOVA: F_(1,34)_ = 0.17, p = 0.67 for ‘treatment’; F_(2,68)_ = 550.8, p = 2×10^−16^ for ‘state’; F_(2,68)_ = 0.59, p = 0.55 for interaction. (**G**) Delta power dynamics across two light and dark phases and after a 6 h-sleep deprivation. SD, Sleep deprivation, Rec, Recovery period, Ctrl control periods with corresponding ZT values. Delta power values are normalized to the mean of those at ZT9-12. Shaded areas represent SEM. SD was carried on a subset of 12 Sham and 12 SNI from the 18 shown in A—F, directly following the D20+ recording. (**H**) Boxplot for delta power values during Rec. One-sample *t*-test for Sham: t_(11)_ = 17.2, p = 2.6×10^−9^; SNI: t_(11)_ = 16.48, p = 4.2×10^−9^; between Sham and SNI two-sample *t*-test: t_(22)_ = − 0.6, p = 0.52; α = 0.0125. (**I, J**) As panels G, H for the number of MAs per h of NREMS. (**J**) Mixed-model ANOVA: F_(1,22)_ = 0.9, p = 0.35 for ‘treatment’; F_(1,22)_ = 58.52, p = 1.2×10^−7^ for ‘SD’; F_(1,22)_ = 0.12, p = 0.72 for interaction. (**K, L**) As panels G, H, for the preferred phase of the 0.02 Hz-fluctuation at MA onset. (**L**) Mixed-model ANOVA: F_(1,22)_ = 3.08, p = 0.09 for ‘treatment’; F_(1,22)_ = 1.45, p = 0.24 for ‘SD’; F_(1,22)_ = 5.72, p = 0.025 for interaction. No significance in paired *post-hoc t*-tests.

It has been shown that sleep loss exacerbates pain (Alexandre *et al*., 2017). Sleep could thus be relatively more disrupted in SNI animals after a period of sleep loss. We therefore carried out a 6 h-sleep deprivation (SD) at the beginning of the light phase as done previously in the lab (n = 12 for Sham and SNI each) (Kopp *et al*., 2006). We confirmed a characteristic rebound of delta power (***Figure 3G***,***H***) and a decrease in the frequency of MAs (***Figure 3I,J***, mixed-model ANOVA with factors ‘treatment’ and ‘SD’, p = 0.35 and p = 1.23×10^−7^ with no interaction). The phase-coupling of MAs to the 0.02 Hz-fluctuation remained unaltered in both groups even with high sleep pressure (***Figure 3K,L***). Conditions of SNI thus left spontaneous MAs, their coupling to the 0.02 Hz-fluctuation, as well as homeostatic regulation of spontaneous arousability unaltered.

### NREMS in SNI conditions shows a novel type of cortical local arousal

In human NREMS, spontaneous arousals are an important measure for the severity of sleep disorders and are primarily described by EEG desynchronization (Bonnet *et al*., 1992; Azarbarzin *et al*., 2014). In contrast, the scoring of a MA in mice requires concomitant muscular activity by convention (Franken et al., 1999). We hence tested whether the 0.02 Hz-fluctuation could serve to identify previously undescribed arousal types in mice with characteristics distinct from conventional MAs. For this, we generated spectral profiles of all cycles of the 0.02 Hz-fluctuation that were devoid of MAs. To take into account the possibility that there were local events delimited to certain cortical regions (Nobili *et al*., 2011; St-Jean et al., 2012; Riedner et al., 2016; Lecci et al., 2017), we combined polysomnography with stereotaxically guided local field potential (LFP) recordings, as done previously in the lab (***Figure 4A,B***) (Fernandez *et al*., 2018). We chose the S1 hindlimb (S1HL, 5 Sham and 9 SNI) cortex (***Figure 4C,D***) that is the site of sensory discrimination of pain and the prelimbic (PrL, 6 Sham and 8 SNI) cortex (***Figure 4I,J***) that is concerned with aversive pain feelings in rodents (Kuner & Kuner, 2020) and in its homologue in humans (Moisset & Bouhassira, 2007).

**Figure 4.**
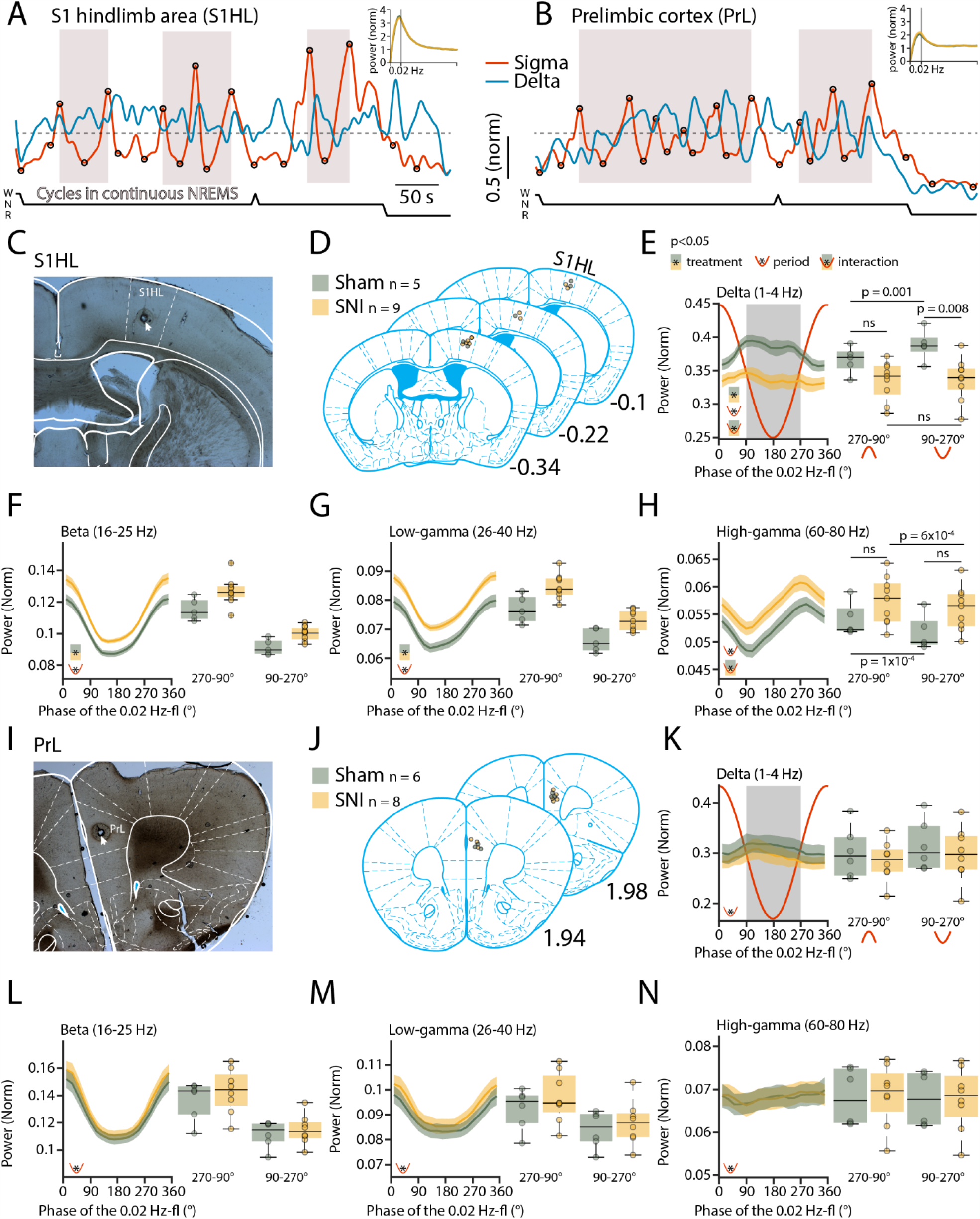
SNI animals present locally disrupted spectral power dynamics during NREMS. (**A-B**) Sigma (10— 15 Hz) and delta (1—4 Hz) dynamics during the same NREMS period in S1 hindlimb (HL) (A) and PrL (B) during a baseline recording. Hypnograms shown below. Circles represent peaks and troughs used to detect the 0.02 Hz-fluctuation cycles (see Methods). Shaded areas indicate 0.02 Hz-fluctuation cycles that are continuing uninterrupted in NREMS. Insets: Power spectrograms in the infraslow range, with Sham and SNI overlapping. (**C**) Representative histological section confirming the location of recording site through post-mortem electrocoagulation (arrowhead). (**D**) Anatomical sections summarizing histological verification, with small circles showing recording sites for all animals included. Anteroposterior stereotaxic coordinates are given relative to Bregma. (**E-H**) Trajectories of power in specific frequency bands (indicated on top of graphs) across uninterrupted cycles of the 0.02 Hz-fluctuation, quantified in 20° bins. Red line shows the corresponding 0.02 Hz-fluctuation phase. Boxplots quantify spectral power within continuity (270—90°, red inverted U-shaped line) and fragility periods (90—270°, red U-shaped line). For all bands, mixed-model ANOVA with factors ‘treatment’ and ‘period’ were done, followed by *post-hoc t*-tests if applicable. Significance for main effects and/or interaction are shown by the presence of the corresponding pictograms at the bottom left. These yielded (**E**) <u>for delta</u> F_(1,12)_ = 7.16, p = 0.02 for ‘treatment’; F_(1,12)_ = 18.5, p = 0.001 for ‘period’; F_(1,12)_ = 18.31, p = 0.001 for interaction. *Post-hoc* for Sham *vs* SNI in continuity: t_(12)_ = 2.11, p = 0.055; fragility: t_(12)_ = 3.13, p = 0.0085; and across periods for Sham: t_(4)_ = −7.9, p = 0.001; SNI: t_(8)_ = −0.8, p = 0.44; α = 0.0125; (**F**) <u>for beta</u> F_(1,12)_ = 9.3, p = 0.01 for ‘treatment’; F_(1,12)_ = 270.01, p = 1.3×10^−9^ for ‘period’; F_(1,12)_ = 1.01, p = 0.33 for interaction. *Post-hoc* for Sham *vs* SNI in continuity: t_(12)_ = −2.5, p = 0.025; fragility: t_(12)_ = −3.43, p = 0.004; and across periods for Sham: t_(4)_ = 14.7, p = 0.0001; SNI: t_(8)_ = 12.02, p = 2.1×10^−6^; α = 0.0125; (**G**) <u>for low-gamma</u> F_(1,12)_ = 11.49, p = 0.0053 for ‘treatment’; F_(1,12)_ = 398.35, p = 1.4×10^−10^ for ‘period’; F_(1,12)_ = 0.75, p = 0.4 for interaction. *Post-hoc* for Sham *vs* SNI in continuity: t_(12)_ = −3.1, p = 0.008; fragility: t_(12)_ = −3.4, p = 0.0049; and across periods for Sham: t_(4)_ = 18.6, p = 4.9×10^−5^; SNI: t_(8)_ = 14.39, p = 5.3×10^−7^; α = 0.0125; (**H**) <u>for high-gamma</u> F_(1,12)_ = 3.31, p = 0.09 for ‘treatment’; F_(1,12)_ = 94.38, p = 4.8×10^−7^ for ‘period’; F_(1,12)_ = 5.83, p = 0.03 for interaction. *Post-hoc* for Sham *vs* SNI in continuity: t_(12)_ = −1.5, p = 0.14; fragility: t_(12)_ = −2.08, p = 0.059; and across periods for Sham: t_(4)_ = 14.17, p = 1.4×10^−4^; SNI: t_(8)_ = 5.45, p = 6×10^−4^; α = 0.0125. (**I-N**) Analogous presentation for PrL. Statistical analysis yielded: (**K**) <u>For delta</u> F_(1,12)_ = 0.4, p = 0.5 for ‘treatment’; F_(1,12)_ = 11.99, p = 0.004 for ‘period’; F_(1,12)_ = 0.009, p = 0.92 for interaction; (**L**) <u>for beta</u> F_(1,12)_ = 0.56, p = 0.46 for ‘treatment’; F_(1,12)_ = 127.5, p = 9.5×10^−8^ for ‘period’; F_(1,12)_ = 0.65, p = 0.43 for interaction; (**M**) <u>for low-gamma</u> F_(1,12)_ = 0.63, p = 0.44 for ‘treatment’; F_(1,12)_ = 75.84, p = 1.5×10^−6^ for ‘period’; F_(1,12)_ = 0.93, p = 0.35 for interaction; (**N**) <u>for high-gamma</u> F_(1,12)_ = 0.004, p = 0.94 for ‘treatment’; F_(1,12)_ = 9.49, p = 0.009 for ‘period’; F_(1,12)_ = 2.6, p = 0.12 for interaction.

Local field potential recordings reliably reported on the 0.02 Hz-fluctuation in these two areas. Consistent with its predominant expression in sensory cortices (Lecci et al., 2017), the 0.02 Hz-fluctuation showed a higher peak in S1HL than in PrL (***Figure 4A,B***). The cycles of successive continuity and fragility periods were extracted (***Figure 2, figure supplement 1***) and their spectral dynamics plotted separately for the relative contribution of power in the low-frequency delta (1—4 Hz) and the beta (16—25 Hz), low-(26—40 Hz) and high-(60—80 Hz) gamma bands (***Figure 4E-H*** for S1HL, ***Figure 4K-N*** for PrL). Average values for the infraslow phase angles between 90—270°, corresponding to the fragility period enriched in MAs (*see* ***Figure 2***), and for the continuity period (from 270—90°), were calculated. Such analysis revealed SNI- and region-specific alterations in the contributions of these bands to total power that were clearly present in S1HL, but not detectable in PrL. In S1HL, delta power levels were decreased compared to Sham (***Figure 4E***) whereas high-frequency components in the beta and the low-gamma range were elevated (***Figure 4F-G***). Remarkably, delta power differences between Sham and SNI varied between fragility and continuity periods (***Figure 4E***, mixed-model ANOVA with factors ‘treatment’ and ‘period’, p = 0.001 for the interaction). In Sham, there was a distinct rapid upstroke of power in this frequency band that reached a peak during the fragility periods (***Figure 4E***, *post-hoc t*-test for delta power in fragility vs continuity period in Sham, p = 0.001). Fragility periods continuing into NREMS were thus clearly distinct from the ones associated with MAs during which there is muscular activity and a decrease in EEG delta power (see ***Figure 2C***). In SNI animals, in contrast to Sham, there was no detectable elevation in delta power during fragility periods continuing into NREMS (***Figure 4E***, *post-hoc t*-test in SNI, p = 0.44). The high-frequency bands in the beta and low-gamma range instead showed a tonic increase in SNI that was present throughout continuity and fragility periods (***Figure 4F-G***, mixed-model ANOVA with factors ‘treatment’ and ‘period’, for beta, p = 0.01, p = 1.3×10^−9^ and for low gamma, p = 0.0053, p = 1.4×10^−10^) and that was also present, although to a milder extent, in the high-gamma range (***Figure 4H***).

We calculated an “activation index” (AI), defined by the ratio between the summed spectral power in the beta and low-gamma bands and the delta band power, to quantify alterations in spectral balance between high- and low-frequency power components, similarly to what has been done previously in studies on insomnia disorders (Lecci et al., 2020). The AI is a measure for the degree of EEG desynchronization and increases when NREMS moves closer to wakefulness. In the fragility periods continuing into NREMS and devoid of EMG activity, the AI decreased, consistent with NREMS remaining consolidated (***Figure 5A-C***). In SNI animals, however, the AI was higher compared to Sham specifically in the fragility periods (***Figure 5B***, mixed-model ANOVA with factors ‘treatment’ and ‘period’, p = 0.039 for interaction, *post-hoc t*-tests Sham vs SNI in fragility period, p = 0.005, in continuity period, p = 0.027, not significant with α = 0.0125). Fragility periods during uninterrupted NREMS are thus specific moments during which the AI in SNI conditions was significantly higher compared to continuity periods.

**Figure 5.**
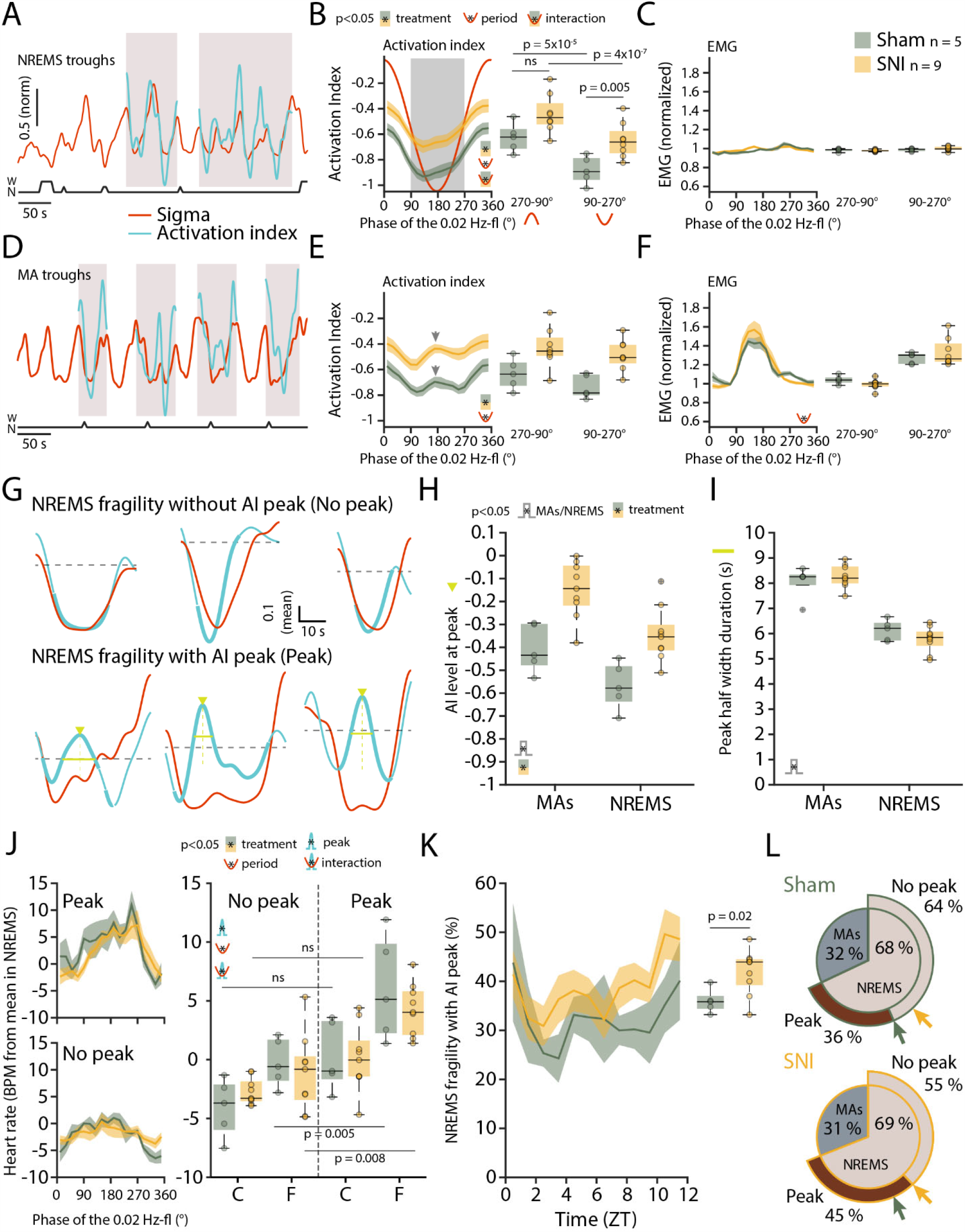
Mice generate local cortical arousals during NREMS that appear more frequently in SNI. (**A**) Normalized dynamics of sigma (10—15 Hz) and activation indices (calculated as ln((beta + low-gamma) /delta))). Hypnogram is shown below. Shaded areas show cycles of the 0.02 Hz-fluctuation continuing into NREMS. (**B**) Activation indices for cycles continuing into NREMS, with mean values shown in boxplots for continuity (270—90°) and fragility periods (90—270°). Mixed-model ANOVA F_(1,12)_ = 8.9, p = 0.011 for ‘treatment’; F_(1,12)_ = 470.6, p = 5.4×10^−11^ for ‘period’; F_(1,12)_ = 5.33, p = 0.039 for interaction. *Post-hoc t*-test for Sham *vs* SNI in continuity: t_(12)_ = −2.51, p = 0.027; fragility: t_(12)_ = −3.44, p = 0.005; and across periods for Sham: t_(4)_ = 17.98, p = 5.6×10^−5^; SNI: t_(8)_ = 14.86, p = 4.1×10^−7^; α = 0.0125. (**C**) Corresponding EMG values (normalized to the mean value in NREMS). Mixed-model ANOVA F_(1,12)_ = 0.2, p = 0.65 for ‘treatment’; F_(1,12)_ = 3.83, p = 0.07 for ‘period’; F_(1,12)_ = 1.31, p = 0.27 for interaction. (**D-F**) Same presentation as in panels A-C, for cycles of the 0.02 Hz-fluctuation associated with MAs. Shaded areas in panel D show cycles of the 0.02 Hz-fluctuation interrupted by a MA. Arrowhead in E denotes the peak of the intermittent increase in AI due to MA occurrence. Statistics for E: Mixed-model ANOVA F_(1,12)_ = 11.26, p = 0.0057 for ‘treatment’; F_(1,12)_ = 25.34, p = 2.9×10^−4^ for ‘period’; F_(1,12)_ = 2.61, p = 0.13 for interaction. Statistics for F: Mixed-model ANOVA F_(1,12)_ = 0.045, p = 0.83 for ‘treatment’; F_(1,12)_ = 44.5, p = 2.3×10^−5^ for ‘period’; F_(1,12)_ = 0.91, p = 0.35 for interaction. (**G**) Six individual cases in one sham animal illustrating sigma (red) and AI (blue) dynamics in uninterrupted cycles of the 0.02 Hz-fluctuation. Thick portions of the blue traces represent the AI during the fragility period. Top three examples show an AI without an intermittent peak, bottom three examples show an AI with a peak (Peak is denoted by green arrowheads and duration at half-maximum by the green line). The horizontal line (mean AI per cycle) represents the threshold for peak detection. (**H**) Values of AI at the intermittent peak for cycles with (left, MA) and without (right, NREMS) an interruption by a MA. Mixed-model ANOVA F_(1,12)_ = 14.94, p = 0.0022 for ‘treatment’; F_(1,12)_ = 110.29, p = 2.1×10^−7^ for ‘MA’; F_(1,12)_ = 0.76, p = 0.39 for interaction. (**I**) Duration of the intermittent peak at half-maximum, for cycles with (left, MA) and without (right, NREMS) an interruption by a MA. Mixed-model ANOVA F_(1,12)_ = 0.24, p = 0.62 for ‘treatment’; F_(1,12)_ = 101.64, p = 3.28×10^−7^ for ‘MA’; F_(1,12)_ = 1.39, p = 0.26 for interaction. (**J**) Left: Heart rate dynamics (compared to the mean heart rate in NREMS), for cycles continuing into NREMS. Cycles are divided in whether a peak in activation index (AI) was present (top) or not (bottom). Right: boxplot quantification of the heart rate values shown on the left for continuity (270—90°) and fragility periods (90—270°), for Sham and SNI in NREMS fragility periods without (left of the dotted line) or with (right of the dotted line) an AI peak. Mixed-model three-way ANOVA F_(1,12)_ = 1.75, p = 0.21 for ‘treatment’; F_(1,12)_ = 26.71, p = 2.3×10^−4^ for ‘peak’; F_(1,12)_ = 11.18, p = 0.005 for ‘period’; only significant interaction in ‘peak’ x ‘period’ F_(1,12)_ = 10.7; p = 0.007. *Post-hoc* cycles with AI peak vs cycles without AI peak for sham in continuity: t_(4)_ = −4.01, p = 0.015; Sham in fragility: t_(4)_ = −4.61, p = 0.009; SNI in continuity: t_(8)_ = −2.82, p = 0.022; SNI in fragility: t_(8)_ = −3.51, p = 0.008; α = 0.0125. (**K**) Occurrence of fragility periods with intermittent peak in AI across the light phase, quantified in boxplots on the right (unpaired *t*-test for Sham *vs* SNI t_(12)_ = −2.66, p = 0.02). (**L**) Two level-pie plots for Sham (left) and SNI (right) representing the proportion of cycles containing a MA or continuing in NREMS. These latter fragility periods are further subdivided into the ones with intermittent peak (‘Peak’) and without intermittent peak (‘No peak’). The arrows show the proportions for Sham (green) and SNI (yellow).

Can such mean differences in cortical activation profiles during NREMS qualify as differences in cortical arousals? To address this, we compared the AI in fragility periods *with* a MA (associated with EMG increase). As expected, the AI showed an intermittent phasic peak (***Figure 5D-F***) in most cases (75.2 ± 4.1 %), which is explained by the strong decline in delta power (see ***Figure 2C***) and the appearance of higher frequencies associated with MAs. Therefore, we inspected individual fragility periods continuing into NREMS (without a MA) for the presence of similar phasic increases in AI. Indeed, we noticed that a subset of these did indeed contain an intermittent peak resembling the one found during MAs (***Figure 5G***) and not evident in the mean AI in ***Figure 5B***. The amplitudes of these peaks were higher in SNI, in accordance with the tonically higher AI in these animals, but in size comparable to the ones of MAs (***Figure 5H***, mixed-model ANOVA with factors ‘treatment’ and ‘MA’, p = 0.002 for ‘treatment’, p = 2.1×10^−7^ for ‘MA’, no interaction). Moreover, the half-widths of these peaks were only moderately smaller than the ones of MAs (***Figure 5I***, mixed-model ANOVA with factors ‘treatment’ and ‘MA’, p = 0.62 for ‘treatment’, p = 3.28×10^−7^ for ‘MA’, no interaction). These events could thus qualify as a local cortical arousal based on phasic spectral properties reminiscent of a MA. To further support our assumption that these AI peaks constituted arousals, we looked at heart rate increases known to accompany cortical arousals in human (Sforza *et al*., 2000; Azarbarzin et al., 2014). The heart rate was distinctly higher during the fragility period for cycles containing an AI peak as opposed to the ones without such peak (**Figure 5J**, mixed-model ANOVA with factors ‘treatment’, ‘period’ and ‘peak’, p = 0.007 for the ‘peak’ x ‘period’ interaction). These events were more frequent in SNI animals and followed a similar time-of-day dependence as the classical MAs (***Figure 5K,L***, *t*-test Sham *vs* SNI, p = 0.02). Moreover, their increased occurrence was specific for S1HL while absent in PrL and in the contralateral EEG (***Figure 5 – figure supplement 2***). The presence of a subgroup of fragility periods continuing into NREMS, yet showing a cortical arousal, is noteworthy for several reasons. First, it demonstrates that rodent NREMS shows local cortical intrusion of wake-related activity in the absence of muscular activity. Second, these local cortical arousals in SNI showed intermittent peaks in AI that were close the ones of MAs, indicating comparable cortical desynchronization at the local level. Third, they were accompanied by heart rate increases that are sensitive hallmarks of arousal in human (Azarbarzin et al., 2014). Fourth, neuropathic pain goes along with a specific increase in the relative occurrence of fragility periods with such AI peaks specifically in the S1HL area. The systematic classification of fragility periods helped unravel these novel cortico-autonomic arousals and their similarity to MAs. Still, other arousal-like events outside fragility periods could exist.

### The 0.02 Hz-fluctuation allows to anticipate elevated spontaneous arousability during NREMS

We finally examined sensory arousability in SNI conditions, focusing on the somatosensory modality. To anticipate fragility and continuity periods in real-time in the sleeping animal, we trained a machine learning software to predict online periods of continuity and fragility based on EEG/EMG recordings (***Figure 6A-E***). For the training, we used online-calculated 0.02 Hz-fluctuation estimates onto which fragility and continuity periods were labelled using peak-and-trough detection of sigma power dynamics (***Figure 6 – figure supplement 3***). To control for the accuracy of the online prediction, we visually scored MAs in 12 C57Bl/6J animals implanted only for polysomnography and verified their position in either online detected peak-to-trough (‘online fragility’) or trough-to-peak phases (‘online continuity’) (***Figure 6F***). We compared the online prediction to that generated by chance through randomly shuffling both online fragility and continuity point positions in the recordings. This showed that the MA proportions obtained with the real detection exceeded those obtained by chance prediction (***Figure 6G***, for online fragility periods, p = 0.0004, for online continuity periods, p = 0.0028). Online detection of peak-to-trough and trough-to-peak periods of the 0.02 Hz-fluctuation is thus a versatile method to probe variations of evoked arousability from NREMS.

**Figure 6.**
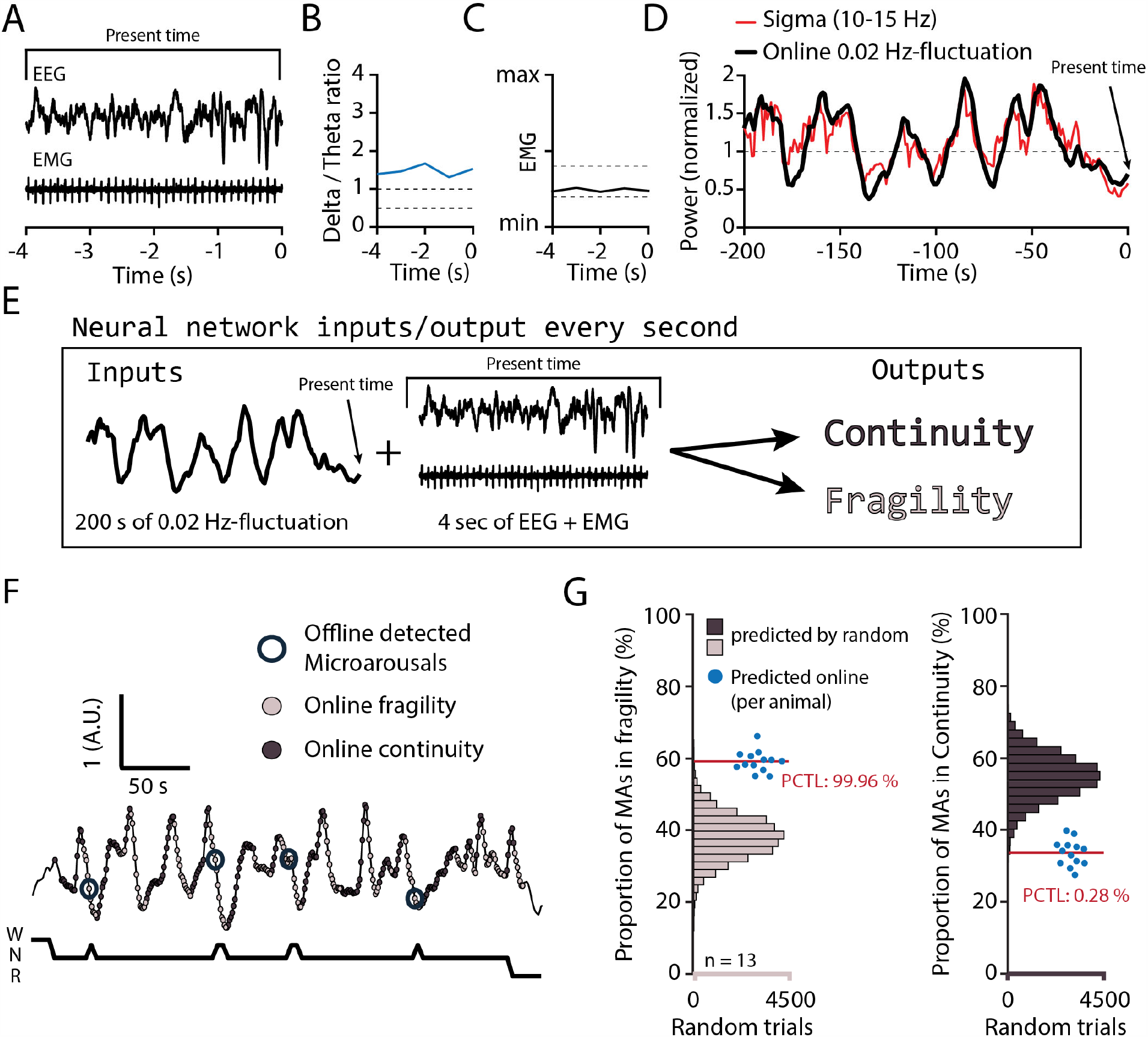
On-line detection of fragility periods during mouse NREMS. (**A-E**) Input and output parameters for machine-learning of NREMS fragility and continuity periods. From the EEG and EMG up to the momentary time. (**A**) 4 last second of EEG and EMG at momentary time. (**B**) Ratio delta (1—4 Hz) over theta (5—10 Hz) from the FFT every s (**C**) normalized EMG levels. Together (B,C), are the variables for state decision using appropriate thresholds (dashed lines, see Methods). (**D**) The 0.02 Hz-fluctuation was estimated (black line) from sigma values (red) every s. The black line was constructed using the last point of a 9^th^ order polynomial fit over the last 200 s of sigma power values. (**E**) The network was trained to use the last 200 s of online 0.02 Hz-fluctuation and the present window of EEG and EMG to determine whether the animal was in continuity or in fragility. The detection was carried out only when the animal was likely in NREMS. (**F**) Representative trace resulting from an online detection of fragility and continuity periods over a period of NREMS. Every s of detection is marked by a dark or light grey circle for online continuity and fragility periods, respectively. The hypnogram below represents the visual scoring done offline, blind to the online detection. The offline-detected MAs are indicated by open circles over the corresponding point of online detection. (**G**) Left, Proportion of MAs scored during online detected fragility. Right, proportion of MAs scored over online detected continuity, for 13 animals (blue dots). Horizontal histograms represent the distribution of possible values of these proportions for randomly shuffled points of fragility or continuity. The mean proportions for the 13 animals fell at percentile 99.96 % for fragility and 0.28 % for continuity.

### SNI conditions produce elevated somatosensory-evoked arousability from NREMS

Evoked arousability was probed through applying sensory stimuli either during online detected fragility or continuity periods. To deliver somatosensory stimuli remotely while the animals were asleep, we attached vibrational motors to their head implant that could be triggered to briefly vibrate (for 3 s) to test the chance for wake-up (***Figure 7A***). These motors were calibrated to vibrate with the same low intensity (∼ 30% of full power) across animals (***Figure 7 – figure supplement 4***). Vibrations were applied randomly with 25% probability during either online detected fragility or continuity periods, for at least two complete light phases per condition (***Figure 7B***). Intensity was chosen such that Sham animals showed approximately equal chances for wake-up or sleep-through in online continuity periods (***Figure 7C***). Moreover, these vibrations produced wake-ups that were short, indicating that the sleeping animal felt only mildly perturbed. Consistent with prior findings, similar stimuli applied during online fragility periods showed consistently higher chances for wake-up (Lecci et al., 2017). In SNI, sensory arousability was elevated for both continuity and fragility periods, leading to highest values during the online fragility periods (***Figure 7D***, mixed-model ANOVA with factors ‘treatment’ and ‘online period’, p = 0.0049 for ‘treatment’, p = 1.31×10^−8^ for ‘online period’, no interaction). Interestingly, consistent with the tonic increase in AI, this increase in sensory arousability in SNI was present across the whole light phase with a conserved time of day dependence (***Figure 7E***).

**Figure 7.**
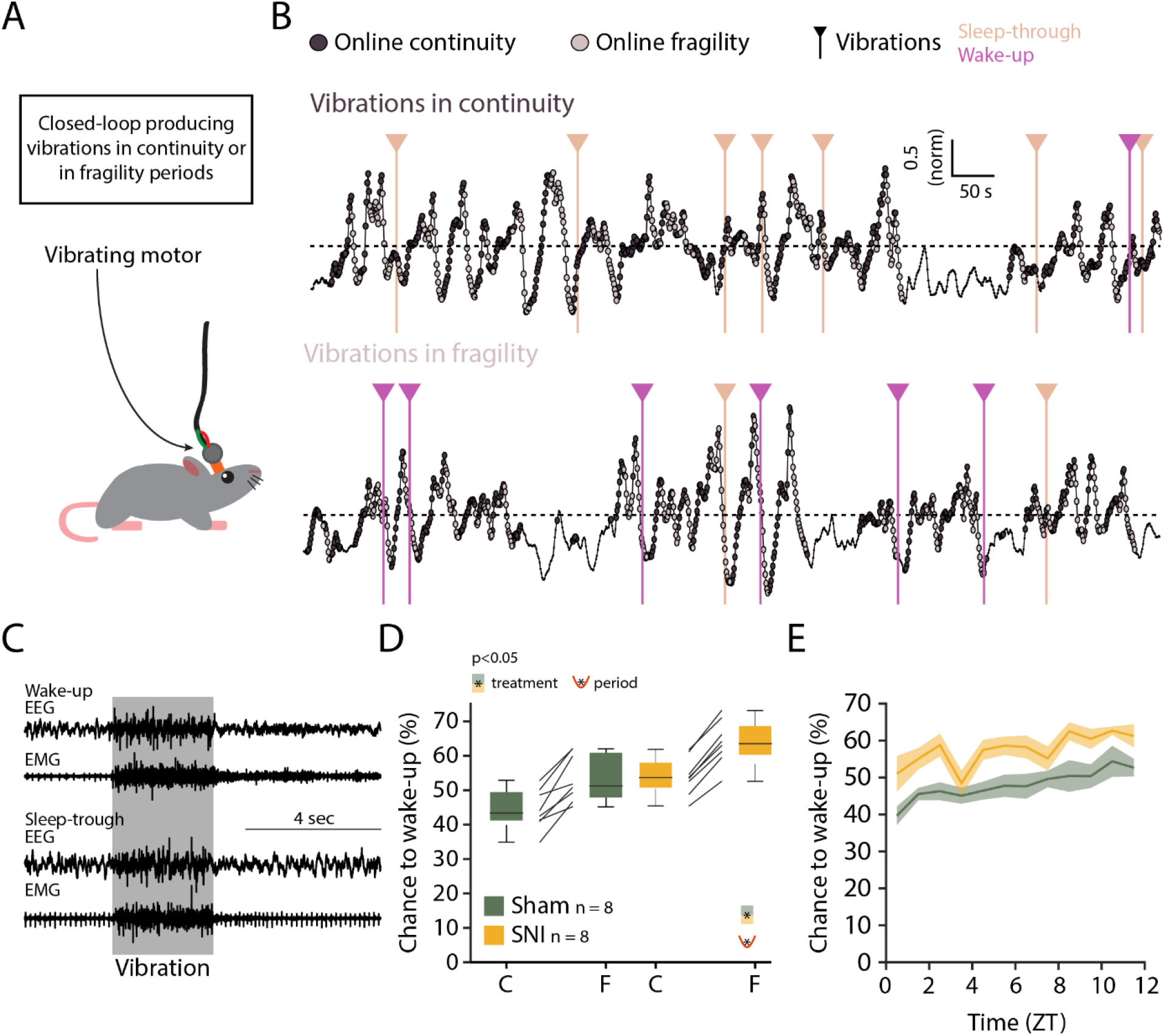
Closed-loop probing of sensory-evoked arousability reveals a more fragile sleep in SNI. (**A**) Setup for delivering vibrating stimuli to the sleeping animal. A small vibrating motor was fixed at the end of the recording cable that was connected to the animal’s headstage. When the algorithm presented in Figure 6 detected a change to an online fragility or continuity for > 4 s, a 3-s mild vibration (see Methods) was delivered with a chance of 25%. (**B**) Example of NREMS periods with vibrations over online continuity (top) or fragility (bottom) periods. Dark and light circles represent the online detection (see Figure 6F). Vertical lines represent the moments when the vibrations were delivered. Color-codes for wake-up or sleep-throughs. (**C**) Representative EEG/EMG recordings representing a wake-up (top) and a sleep-through (bottom). Time of stimulus delivery is shaded. The artifacts caused by the vibrations were not considered during automated wake-up or sleep-through detection (see Methods). (**D**) Chances to wake-up in response to a vibration in online continuity (C) or online fragility (F) periods. Mixed-model ANOVA F_(1,14)_ = 11.07, p = 0.0049 for ‘treatment’; F_(1,14)_ = 136.58, p = 1.31×10^−8^ for ‘period’; F_(1,14)_ = 0.3, p = 0.58 for interaction. (**E**) Probability of wake-up (pooled continuity and fragility) for Sham and SNI over the light phase (ZT0-12). Shaded areas are SEMs.

## Discussion

Chronic pain is a widespread and complex condition compromising sleep. As poor sleep further aggravates pain, therapeutic approaches to improve sleep quality have potential to attenuate disease progression. Here, in efforts to tease apart pain-sleep associations, we decided to focus on mechanisms of sleep disruptions at early stages of chronic pain. This study progresses on the sleep-pain association in four essential ways. First, we show that brain and autonomic signatures of pain states during the day intrude in a persistent manner into sleep in early phases of the disease. Second, sleep appeared nevertheless preserved in architecture, in dominant spectral band power, and in homeostatic regulation. Third, a previously undescribed spontaneous arousal during NREMS, showing local cortical activation with concomitant heart rate increases, appeared more frequently in SNI conditions. Fourth, we also demonstrate that fine mechanovibrational stimuli triggered brief wake-ups from NREMS more easily in SNI animals. In summary, chronic pain impacts on NREMS in terms of arousability, more specifically in the probability that NREMS transits towards diverse levels of wakefulness, either spontaneously, or with external stimuli. Chronic pain additionally instates a tonic regional elevation of high-frequency electrical activity. Both these consequences are not detectable in conventional measures of sleep. However, they bear resemblance to pathophysiological markers of insomnia disorders and, as we show here, produce elevated responsiveness to fine vibrational stimuli. The study further suggests that, amongst the many peripheral and central circuit alterations caused by chronic pain, the ones affecting the primary nociceptive sensory areas could serve as a site of entry to treat pain-related sleep disturbances directly at early states of the disease. For example, targeted interference by transcranial stimulation techniques has been proposed to modulate pain-related oscillations and could thus be probed to modify abnormal arousals (Shirvalkar *et al*., 2018; Hohn *et al*., 2019).

The quantification and classification of arousals from NREMS, both in terms of their physiological correlates and in their intensity, is central to estimate the severity of a sleep disorder. The fragmentation of sleep by arousals is the primary cause for daytime fatigue and for cognitive deficits and, in more severe cases, may constitute long-term risks for cardiovascular health (Bonnet et al., 1992; Silvani, 2019). Criteria for arousal scoring in humans are comparatively well established and there are strong indications that arousal intensity is graded, with EEG desynchronization and concomitant heart rate acceleration occurring independently of muscular activity (e.g. leg movements) (Sforza et al., 2000; Azarbarzin et al., 2014). In chronic pain patients, few systematic analyses on spontaneous arousals are currently available and a need for more polysomnographic assessments in well-controlled patient populations has been highlighted (Bjurstrom & Irwin, 2016). In rodents, only few studies have described cortical desynchronization events without EMG activity, and these were not characterized with respect to autonomic correlates (Bergmann *et al*., 1987; Franken, 2002; Léna *et al*., 2004; Fulda, 2011). This relative lack of arousal characterization in mouse NREMS could have hampered the identification of models to replicate sleep disorders, as we found here to be the case for chronic pain models. To the best of our knowledge, our analysis of the SNI mouse is the first that qualifies as a rodent model replicating physiological correlates of insomnia disorders that are hidden behind a comparatively normal sleep and that could raise awareness for refined analysis of the diverse forms of sleep disruptions in chronic pain patients (Bjurstrom & Irwin, 2016). Our work also adds a novel variant to proposed insomnia models that provoked severe macroscopic sleep disruptions either through acute stress (Cano *et al*., 2008; Li *et al*., 2020) or by optogenetically enforced full awakenings to fragment NREMS (Rolls *et al*., 2011).

This study proposes a novel type of arousal from NREMS in mouse that pairs cortical desynchronization with heart rate increases. Inclusion of this type of arousal is crucial to identify the exact sleep disruptions for the case of neuropathic pain. We departed here from previous observations on infraslow variations in sensory arousability during NREMS that take place over the minute time-scale (Lecci et al., 2017). We first demonstrated that spontaneous MAs, the only spontaneous arousal in rodent for which scoring criteria are widely established, occur remarkably clustered at phases for which sensory evoked arousals were most likely (Lecci et al., 2017). The high number of MAs and their remarkable clustering at phases > 90° and < 270°, allowed us to allocate the fragility periods to the phases with low sigma power. We consider this basic finding on MA timing in rodent NREMS significant for several reasons. First, it suggests that the infraslow 0.02 Hz-fluctuation is part of an overarching process that periodically sets a fragility of NREMS towards wake-promoting inputs, whether they arise from internal processes or external stimuli. Mechanistically, this points to an involvement of widely projecting neuromodulatory brain areas such as the locus coeruleus that remains active during NREMS (Aston-Jones & Bloom, 1981; Kjaerby *et al*., 2020) and regulates sensory arousability (Hayat *et al*., 2020). Second, the finding contributes to a long-standing uncertainty about the origin and the stochastic time-scales on which MAs are thought to occur (Lo *et al*., 2004; Dvir *et al*., 2018). We now calculate that MAs in consolidated NREMS occur with a mean ∼35% probability on ∼50-sec intervals, thus providing a temporal raster on which these can be anticipated with a measurable degree of certainty. MAs depend on activity in wake-promoting brain areas (Dvir et al., 2018), specifically on the histaminergic hypothalamus in mice (Huang *et al*., 2006) and on cholinergic nicotinic receptors (Léna et al., 2004). This mechanistic origin of MAs is consistent with our observation that they occur at moments of NREMS during which sensory wake-ups occur preferentially. We therefore consider the identification of fragility periods as time raster for MAs as key to reinforce mechanistic investigations into the origins of spontaneous arousals. This not only concerns MAs, but opens the opportunity to search for other arousal-like events that could be relevant to model pathological conditions of human patients.

The majority of this study was dedicated to providing a proof-of-concept for the usefulness of the infraslow fragility periods to scrutinize arousability. The chosen SNI model appeared particularly appropriate for this purpose because it produced a sensory deficit that could be exploited to specifically test somatosensory arousability. The separation of NREMS into fragility and continuity periods throughout the resting phase allowed us to sample and scrutinize the many fragility periods continuing apparently uninterrupted into NREMS. The fragility periods were also critical to determine when AI became most disparate between SNI and Sham and to identify previously undescribed cortico-autonomic arousals. During these, the activation index increased because of a phasic decrease in low-frequency delta power and an increase in high-frequency power. These events were more pronounced and more frequent in SNI because both these phasic power alterations were disrupted, with the deficits in delta power most pronounced. Without the raster provided by the fragility periods, the phasic differences amidst the tonically elevated high-frequency power would easily have gone undetected. To further ascertain that these detected events nevertheless constituted true arousals rather than accidental spectral fluctuations, we sought for independent physiological correlates. Inspired by the human literature (Sforza et al., 2000; Azarbarzin et al., 2014), we found that heart rate increases were consistently higher when calculated for cortical arousals with AI peak than for the ones without AI peak. Moreover, their distribution across the resting phase was similar to the one found for MAs and they were present in both S1HL and PrL. This result supports our interpretation that we have identified here a novel cortical-autonomic arousal subject to similar time-of-day-dependent regulatory mechanisms. Still, we cannot exclude that arousal subtypes outside the fragility periods went undetected that would require further characterization. We also remark here clearly that, aside from more frequent cortico-autonomic arousals, SNI animals suffered from a tonically elevated high-frequency power in S1HL that likely underlay the more elevated sensory arousability throughout continuity and fragility periods.

The lack of major sleep disruptions in the SNI mouse model was initially unexpected but seemed in line with other studies. We analyzed these animals at a time point when pain from the wound and associated inflammations are largely over (Guida *et al*., 2020), both of which can strongly disrupt sleep (Landis *et al*., 1989; Andersen & Tufik, 2003; Silva *et al*., 2008). Moreover, the animals showed a preserved time spent in REMS, suggesting that they did not suffer from chronic mild stress-inflicted sleep disruptions (Nollet *et al*., 2019). Other studies on chronic pain also report diverse moderate effects on sleep (Kontinen et al., 2003; Leys et al., 2013). One study on rats at 2 and 10 days after SNI surgery suggested that brain states intermediate between NREMS and wakefulness during the resting phase exist (Cardoso-Cruz et al., 2011), which could in part reflect our observations. We also found no alterations in theta power or for shifts in theta peaks in wakefulness, as reported for other animal models of chronic pain (LeBlanc *et al*., 2014) or for humans with severe neurogenic pain or arthritis (Sarnthein *et al*., 2006). Analysis of sleep disruptions at later stages in the disease will help decide whether distinct phases of sleep disruptions mark distinct phases of pain chronicity when anxiety- and depression-related behaviors appear more strongly (Guida et al., 2020).

The spectral dynamics in two cortical regions we present here delineate possible areas of pathological neuronal activity that underlie the cortical arousals. The continued presence of high-frequency activity in S1HL is reminiscent of the cortical oscillatory activity evoked with acute painful stimuli, suggesting that nociceptive input continues to arrive in cortex during NREMS to generate excessive excitation. Indeed, it has been suggested that the SNI model does show spontaneous ectopic electrical activity in peripheral sensory neurons as a result of nerve injury (Wall & Devor, 1983; Devor, 2009). NREMS is thought to protect relatively weakly from nociceptive inputs (Claude *et al*., 2015), therefore possibly allowing continued processing of spontaneous nociceptive activity that could explain the cortical spectral changes we detected. It has also been shown that optogenetic stimulation of the thalamic reticular nucleus, known to be implied in the balanced occurrence of delta and spindle waves during NREMS (Fernandez et al., 2018), can alleviate pain in SNI (LeBlanc et al., 2014). Suppressed TRN activity during NREMS could be implied in the attenuated delta dynamics observed in NREMS of SNI mice. In contrast, we found unperturbed local spectral dynamics in PrL during NREMS, although this area is concerned with signaling emotional discomfort in several forms of chronic pain in humans (Schulz *et al*., 2015; Nickel *et al*., 2017; May *et al*., 2019) and is known to undergo strengthened synaptic inhibition in SNI (Zhang *et al*., 2015; Radzicki *et al*., 2017). Further cellular studies will be necessary to understand why these alterations seem not to perturb oscillatory activity in this area during NREMS.

Do SNI animals suffer from insomnia? Our objective measures of NREMS’s spectral composition point to regionally restricted but tonic imbalances in the contribution of low-vs higher frequencies. Patients with insomnia show such imbalances over widespread brain regions that include sensorimotor areas (Lecci et al., 2020). Furthermore, higher power in the beta frequencies has been related to the patients remaining hypervigilant or excessively ruminating at sleep onset (Perlis *et al*., 2001a), preventing the deactivation of cortical processes required for the loss of consciousness. Although insomnia also needs subjective assessments that are not possible in animals, this phenomenological comparison suggests that SNI might suffer from similar experiences due to the tonically enhanced high-frequency oscillations. This interpretation is supported by the elevated wake-up rates in response to mild vibrational stimuli throughout the infraslow cycles, suggesting hyperalertness to environmental disturbance. On top of these tonic changes, there were more frequent cortico-autonomic arousals. Although these do not seem to elevate daytime sleepiness based on the mostly unchanged delta power dynamics across time-of-day, frequent increases in heart rate during the night could augment cardiovascular risk in the long-term (Silvani, 2019). To further analyze the animal’s conditions during daytime, tests on their cognitive abilities in memory-dependent tasks while locally manipulating sleep in the affected hindlimb area could be considered. Deficits in working and declarative memories in rodents with SNI have been documented from early periods of chronic pain (Guida et al., 2020). Chemogenetic manipulation of neuronal populations proposed to be responsible for the gamma activity in chronic pain, restricted to sleep periods (Tan et al., 2019), seems a feasible approach to specifically suppress abnormal pain-related activity during sleep while testing performance in such tasks during wakefulness.

We provide here novel approaches to classify arousals in mouse NREMS that will help in the examination and validation of future candidates for rodent models of sleep disorders. We noted a remarkable stability of the 0.02 Hz-fluctuation across the resting phase that provided us with a temporal raster to screen the characteristics of fragility periods. These led us to identify a previously undescribed cortico-autonomic arousal in mouse NREMS that we also found more frequently in a chronic pain model. Together, this study presents NREMS as a state that is interwoven with arousals showing diverse combinations of physiological parameters with different graduations in intensity that can be the target of pathophysiological changes. Recognizing NREMS as a fluctuating state between fragility and continuity will thus further heighten awareness to arousability as a core component of sleep quality. In this study, we unraveled a so far undescribed sleep disruption in chronic pain that we hope will facilitate further research into the treatment of this devastating condition.

## Materials and methods

### Animal housing and experimental groups

Mice from the C57BL/6J line were singly housed in a temperature- and humidity-controlled environment with a 12-h/12 h light-dark cycle (lights on at 9:00 am, corresponding to ZT0), with access to food and water *ad libitum*. We first used 36 mice, 10-14 weeks-old and bred in our colonies in a conventional-clean animal house, for polysomnography (combined EEG (ECoG)/EMG electrodes), followed by SNI or Sham surgery (18 animals per Sham or SNI group). Mice were transferred from the animal house into the recording room 2-3 d before surgery for polysomnography recording. We recorded a 48 h-long baseline before SNI or Sham surgeries, followed by recording at 22-23 d after surgery (D20+). These data were used for Figures 1 and 3. Total sleep deprivation in Figure 3 was done on 24 of these 36 animals (12 SNI, 12 Sham) within one day following the recording at D20+. The baseline data for Figure 2 were obtained from the baseline recordings of 23 randomly selected animals from the previous 36, completed with 7 more animals from previous baseline recording in the lab. For EEG (ECoG)/EMG/LFP recordings, 33 C57BL/6J male mice of the same age were first operated for SNI or Sham (17 and 14, respectively) and 5 d later, implanted for recordings from S1HL (4 Sham, 6 SNI) or PrL (3 Sham, 4 SNI) or both (3 Sham, 4 SNI). The misplaced or non-functional electrodes were excluded. Recordings were carried on from day 20 to 35 after SNI or Sham surgery. These data were used for Figures 4 and 5. The data of 13 animals previously recorded in the lab and otherwise not included in any dataset in this study were used to train the neural network (EEG/EMG implantation, in *Figure 6*). The experiments on sensory evoked arousals (*Figure 7*) were done on 16 animals (8 Sham, 8 SNI) out of which some (4 sham, 6 SNI) were used for Figures 4-5. All experimental procedures complied with the Swiss National Institutional Guidelines on Animal Experimentation and were approved by the Swiss Cantonal Veterinary Office Committee for Animal Experimentation.

### Surgery for the SNI model of neuropathic pain

The Sham and SNI surgeries were performed as previously described (Decosterd & Woolf, 2000). Briefly, mice were kept under gas anesthesia (1—2 % isofluorane, mixed with O_2_). The left hindleg was shaved and the skin incised. The muscles were minimally cut until the sciatic nerve was exposed. Just below the trifurcation between common peroneal, tibial and sural branches of the nerve, the common peroneal and tibial branches were ligated and transected. The Sham animals, as controls, went through the same surgery without the transection. The muscle and the skin were then stitched closed and the animals were monitored via a score sheet established with the Veterinarian Authorities.

### Surgery for polysomnographic and LFP recordings in mice

Surgeries were performed as recently described (Lecci et al., 2017; Fernandez et al., 2018). Animals were maintained under gas anesthesia (1—2 % isofluorane, mixed with O_2_). Small craniotomies were performed in frontal and parietal areas over the right hemisphere and 2 gold-plated screws (1.1 mm diameter at their base) (Mang & Franken, 2012) were gently inserted to serve as EEG electrodes. Careful scratching of the skull surface with a blade strengthened the attachment of the implant by the glue, so that additional stabilization screws were no longer necessary. Two gold wires were inserted into the neck muscle to serve as EMG electrodes. In the case of LFP recordings, small craniotomies (0.2-0.3 mm) were performed to implant high-impedance tungsten LFP microelectrodes (10–12 MΩ, 75-μm shaft diameter, FHC, Bowdoin, ME) at the following stereotaxic coordinates relative to Bregma in mm, for S1HL: anteroposterior −0.7, lateral −1.8, depth from surface −0.45; for PrL: +1.8, −0.3, −1.45). For the neutral reference for the LFP recordings, a silver wire (Harvard Apparatus, Holliston, MA) was placed in contact with the bone within a small grove drilled above the cerebellum. The electrodes were then soldered to a female connector and the whole implant was covered with glue and dental cement. The animals were allowed 5 d of recovery, while being monitored via a score sheet established with the Veterinarian Authorities, with access to paracetamol (2mg/mL, drinking water). The paracetamol was removed when the animals were tethered to the recording cable for another 5 d of habituation prior to the recording.

### Polysomnographic recording

For sleep recordings, recording cables were connected to amplifier boards that were in turn connected to a RHD USB interface board (C3100) using SPI cables (RHD recording system, Intan Technologies, Los Angeles, CA). For EEG/EMG and/or LFP electrodes, signals were recorded through homemade adapters connected to RHD2216 16-channel amplifier chips with bipolar input or RHD2132 32-channel amplifier chips with unipolar inputs and common reference, respectively. Data were acquired at 1000 Hz via a homemade Matlab recording software using the Intan Matlab toolbox. Each recording was then visually scored in 4-s epochs into wake, NREMS, REMS, as described (Lecci et al., 2017) using a homemade Matlab scoring software.

### Total sleep deprivation protocol

Total SD was carried out from ZT0—ZT6 using the gentle handling method used previously in the lab (Kopp et al., 2006), while animals remained tethered in their home cage. At ZT3, the cages were changed and, from ZT5 to ZT6, new bedding material was provided. At ZT6, the animals were left undisturbed. The recordings carried out during SD were visually scored to assure the absence of NREMS from ZT0 to ZT6. There were no detected NREMS epochs during SD in the mice included in the analysis.

### Probing sensory arousability with vibration motors and automatic wake-up classification

An online detection of continuity and fragility period (described below) was used in a closed-loop manner to time vibration stimuli during NREMS such that sensory arousability could be probed. Small vibrating motors (DC 3—4.2 V Button Type Vibration Motor, diameter 11 mm, thickness 3 mm) were fixed using double-sided tape, at the end of the recording cables, close to the animals’ heads. The motors were driven using a Raspberry Pi 3B+ through a 3.3 V pulse-width modulation (PWM) signal. Each motor was calibrated to find the necessary PWM duty-cycle to output the same amount of mild vibration using a homemade vibration measurer equipped with a piezo sensor. A Python script was running on the Raspberry Pi to detect the voltage change sent by the digital-out channels on the Intan RHD USB interface board. Upon detecting a change from low to high, the Python script waited for an additional 4 s, and assessed the voltage again. In case the voltage was still high, it launched a 3 s vibration with 25% probability. To close the loop, the PWM signals from the Raspberry Pi driving the motors were as well fed into the analog-in channels of the Intan RHD USB interface board to detect the stimuli time-locked to the EEG/EMG signals. In the experiments, the voltage values were set to high during either continuity or fragility, using online detection as described in Data analysis. Four animals could be tested in parallel for their sensory arousability.

### Histological verification of recording sites

After the *in vivo* LFP recordings, the animals were deeply anesthetized with pentobarbital (80 mg/kg) and electrode positions were marked through electro-coagulation (50 µA, 8—10 s). The animals were then transcardially perfused with 4 % paraformaldehyde (in 0.1 M phosphate buffer). After brain extraction and post-fixation for 24 h, 100 µm-thick coronal brain sections were cut and imaged in brightfield microscopy to verify correct electrode positioning.

### Data analysis

#### Scoring and basic sleep measures

Scoring was done blind to the animal treatment according to standard scoring procedures (Fernandez et al., 2018). A MA was scored whenever the EEG presented a desynchronization time-matched with a burst of EMG activity lasting maximally 4 consecutive epochs (16 s). Latency to sleep onset was defined from ZT0 to the first appearance of 6 consecutive NREMS epochs (24 s). The bout size binning in short, intermediate and long bouts for NREMS and REMS was obtained from the pulled distribution of the bout sizes from all the animals. The edges of the intermediate bin were defined as: mean -½ standard deviation to mean + 1 standard deviation.

Spectral power was computed on the raw EEG signal using a FFT on scored 4-s windows after offset correction through subtraction of the mean value of each epoch. The median power spectrum for each state was obtained for epochs non-adjacent to state transitions. The normalization was done through dividing by the average of mean power levels (from 0.75–47 Hz) for each vigilance state, ensuring that each state had the same weight in the averaging (Vassalli & Franken, 2017). This normalization was done separately for Baseline and D20+ recordings. Gamma power at D20+ was extracted through calculating mean power levels between 60—80 Hz. Data were normalized to corresponding baseline values.

The heart rate was extracted from the EMG signal as described previously (Lecci et al., 2017). Briefly, the EMG signal was highpass-filtered (>25 Hz) and squared. The R peaks of the heartbeats were detected using the Matlab ‘Findpeaks’ function. Only animals with clearly visible R peaks present in the EMG in NREMS were included in this analysis (Fernandez *et al*., 2017).

For delta power time course, raw delta power (mean power between 1—4 Hz from FFT on mean-centered epochs) was extracted for each NREMS epoch non-adjacent to a state transition. Total NREMS time was divided into periods of equal amounts of NREMS (12 in light phases, 6 in dark phases) from which mean values for delta power were computed. The position in time of these periods was not different between groups. Normalization was done via mean values between ZT9—12, when sleep pressure is the lowest.

Wake-up and sleep-through events after vibration were scored automatically as follows. For each trial, the EEG and EMG signals were analyzed within time intervals from 5 s prior to 5 s after the vibrations. To distinguish wake-up and sleep-through events, three values were calculated: 1) The ratio theta (5-10 Hz)/ delta (1-4 Hz) for the 5 s before stimulation, 2) the difference in the low-/ high-frequency ratios (1-4 Hz/ 100—500 Hz), before and after the stimulation, 3) the squared EMG amplitude ratio after/ before stimulation.

A trial was rejected when the ratio theta/delta was > 1 before stimulation or the EMG amplitude was larger before than after stimulation. In this way, trials starting in REMS or wakefulness were excluded. Wake-up events were scored when the difference in low/high ratios mentioned above decreased markedly after stimulation together with EMG activity. Occasionally, some wake-ups were also scored when EEG or EMG activity was very high while the other channel showed moderate changes. Appropriate thresholds were set upon visual inspection blinded to the animal’s condition.

#### Analyses related to the 0.02 Hz-fluctuation

##### Extraction

The 0.02 Hz-fluctuation in sigma power (10–15 Hz) was extracted from EEG or LFP signals using a wavelet transform (Morlet wavelet, 4 cycles), calculated over 12 h recordings in 0.5-Hz bins. The resulting signal was down-sampled to 10 Hz and smoothed using an attenuating FIR filter (cutoff frequency 0.0125 Hz, order of 100, the low order allowing for frequencies above the cutoff). The mean of the datapoints within NREMS and MA epochs was used for normalization (***Figure 2-figure supplement 1D***). The peak and frequency of the 0.02 Hz-fluctuation were calculated through a FFT on continuous NREMS bouts as described (Lecci et al., 2017). FFTs from individual bouts at frequency bins from 0 to 0.5 Hz were interpolated to 201 points before averaging across bouts to obtain a single measure per mouse. The angles of the phase of the 0.02 Hz-fluctuation were obtained through the Hilbert transform (Matlab signal processing toolbox). We set the troughs of the 0.02 Hz-fluctuation at 180°, the peaks at 0° (***Figure 2-figure supplement 1G***).

In several instances (**Figures 3D, 4, 5**), instead of calculating FFTs in the infraslow frequency range, we needed to detect individual cycles of the 0.02 Hz-fluctuation. To do this, we applied the Matlab “Findpeaks” function, with the conditions that the peak values were > mean and the trough values < mean, each separated by > 20 s. With such parameters, the sequence trough-peak-trough appears only in NREMS and allows to count individual cycles.

##### Band-limited power dynamics during the 0.02 Hz-fluctuation

To calculate the power dynamics in different frequency bands, Morlet wavelet transforms were down-sampled to 10 Hz to match the sampling of the 0.02 Hz-fluctuation and normalized by the sum of their means in NREMS. The mean power of each band was then binned in 18 bins of 20° and a mean across cycles (with or without MAs) of power activity per phase bin was obtained per animal.

##### Analysis of activation index

AI was computed by the natural logarithm (ln) of the ratio between beta (16– 25 Hz) + low gamma (26–40 Hz) over delta power (1–4 Hz), extracted as described above. Individual cycles from peak-to-peak were classified whether a MA was present in the fragility period or whether it continued into NREMS. To assess the presence of peaks in activation indices, the “findpeaks” function was used at phase values of 90–270°, with mean values used as a threshold.

##### Online detection of continuity and fragility periods

For the online detection of fragility and continuity periods during closed-loop sensory stimulation, a homemade software was generated with two layers of decision. The first one determined the likely current state of vigilance (wake, NREMS or REMS), whereas the second one made a machine learning-based decision between a continuity or a fragility period.

1 – Determination of vigilance state. This assessment was based on power band ratios characteristic for wake, NREMS and REMS using appropriate thresholds (***Figure 6B***,***C***). Every s, a FFT was calculated on the mean-centered last 4 s of EEG values and the power ratio between the delta (1–4 Hz) and the theta (5– 10 Hz) was calculated.

###### Transitions out of wake

*1)* Switch to NREMS if the last 3 s of EMG were below a high threshold and at least 2 out of the 3 last s of ratio were above a high threshold. *2)* Switch to REMS if the full 5 s of EMG were below a low threshold and the full 5 s of ratio were above a high threshold.

###### Transitions out of NREMS

*1)* Switch to wake if the last s of EMG was above a high threshold. *2)* Switch to REMS if the last five s of EMG were below a low threshold and if among the last 5 s of ratio, at least 4 were below a low threshold and all 5 were below a high threshold.

###### Transitions out of REMS

*1)* Switch to wake if the last s of EMG were above a high threshold. *2)* Switch to NREMS if the ratio was above a high threshold for at least 4 out of 5 s.

2 – Continuity and fragility detection: From the previous step, the value of sigma (10—15 Hz) was kept every second. The mean sigma value in NREMS was dynamically updated if the likely state was determined as NREMS and used to normalize the incoming sigma power values. The last 200 s of sigma power regardless of the likely state were kept in memory. We heuristically found that a 9^th^-order polynomial fit (Matlab ‘polyfit’ and ‘polyval’ functions) best approximated the 0.02 Hz-fluctuation. To train the network, we first generated online-estimated 0.02 Hz-fluctuation at 1 Hz for the 12 h of the light phase. We next applied offline cycle detection in NREMS periods. For simplicity, and in agreement with previous measures of sensory arousability (Lecci et al., 2017), we set the continuity periods from trough to peak and fragility from peak to trough. Then, we subdivided these recordings in chunks of 200 s (moving window of 1 s, as they would appear online) and keeping the label continuity, fragility or none for each of them. We could thus obtain 43,000 labelled chunks per 12 h of recording. We used 642,000 of these chunks from 13 animals to train a neural network (pattern recognition ‘nprtool’ from Matlab Statistics and Machine Learning Toolbox) 70 % of the for training, 15 % for validation and 15 % for testing. The network was composed of one hidden layer with 10 neurons and one output layer with the 3 different output. We then used the generated neural network online to take the decision between continuity, fragility or none.

### Statistics

The statistics were done using Matlab R2018a and the R statistical language version 3.6.1. The normality and homogeneity of the variances (homoscedasticity) were assessed using the Shapiro-Wilk and the Bartlett tests, respectively to decide for parametric statistics. In the cases where normality or homoscedasticity were violated, a log transformation was assessed at first and finally, non-parametric *post-hoc* tests were used (Wilcoxon rank sum test for unpaired and signed-rank test for paired data). The degrees of freedom and residuals for the F values are reported according to the R output. *Post-hoc* analyses were done only when the interaction between factors were significant (p < 0.05). Bonferroni’s correction for multiple comparisons was applied routinely, and the corrected α values are given in the legends. The factors used in the ANOVAs are depicted with pictograms once the corresponding effects were significant. The factors used in the analysis were: ‘treatment’ with two levels: Sham and SNI; ‘day’ with two repeated levels: baseline and D20+; ‘size’ with three repeated levels: small, intermediate or long bouts; ‘period’ with two repeated levels: continuity or fragility; ‘SD’ with two repeated levels: control or recovery after sleep deprivation; ‘state’ with three repeated levels: wake, NREMS or REMS; ‘MAs’ with two levels: with or without MA in the fragility period; ‘peak’ with two repeated levels: cycles with or without a peak in AI during fragility periods. The circular statistics were done using the CircStat for Matlab toolbox (Berens, 2009).

## Acknowledgements

All lab members provided critical input at all stages of this manuscript. The excellent animal caretaking headed by Michelle Blom and the Team of Animaliers, in particular Titouan Tromme, is highly appreciated. Expert veterinary support and advice was provided by Drs. Gisèle Ferrand and Laure Sériot. We thank Christiane Devenoges for support in histological analysis and Marie Pertin and Guylène Kirschmann for excellent technical support with SNI surgeries. Dr. Simone Astori and Dr. Marc Suter provided insightful comments on pre-final versions of the manuscript and Laura Solanelles Farré helped with careful proofreading. The useful discussions with Raquel Sandoval Adaia, Paul Franken, Thomas Nevian, Francesca Siclari and Raphaelle Winsky-Sommerer are gratefully acknowledged. This study was funded by The Swiss National Science Foundation (n° 310030_184759 to AL, n° 310030_179169 to ID, n° 320030-179194 to SF), and Etat de Vaud.

## Author contributions

Romain Cardis, Conceptualization, Data collection, Data curation, Formal analysis, Software, Validation, Investigation, Visualization, Methodology, Figures, Writing—review and editing; Sandro Lecci, Conceptualization, Data collection, Data curation, Formal analysis, Methodology; Laura Fernandez, Methodology, Data curation; Alejandro Osorio-Forero, Methodology; Paul Chu Sin Chung, Editing; Stephany Fulda, Conceptualization, Data Validation, Writing—review and editing; Isabelle Decosterd and Anita Lüthi, Conceptualization, Data curation, Supervision, Funding acquisition, Validation, Visualization, Writing—original draft, Project administration, Writing—review and editing.

## Supplementary figures

**Figure 2 – figure supplement 1.**
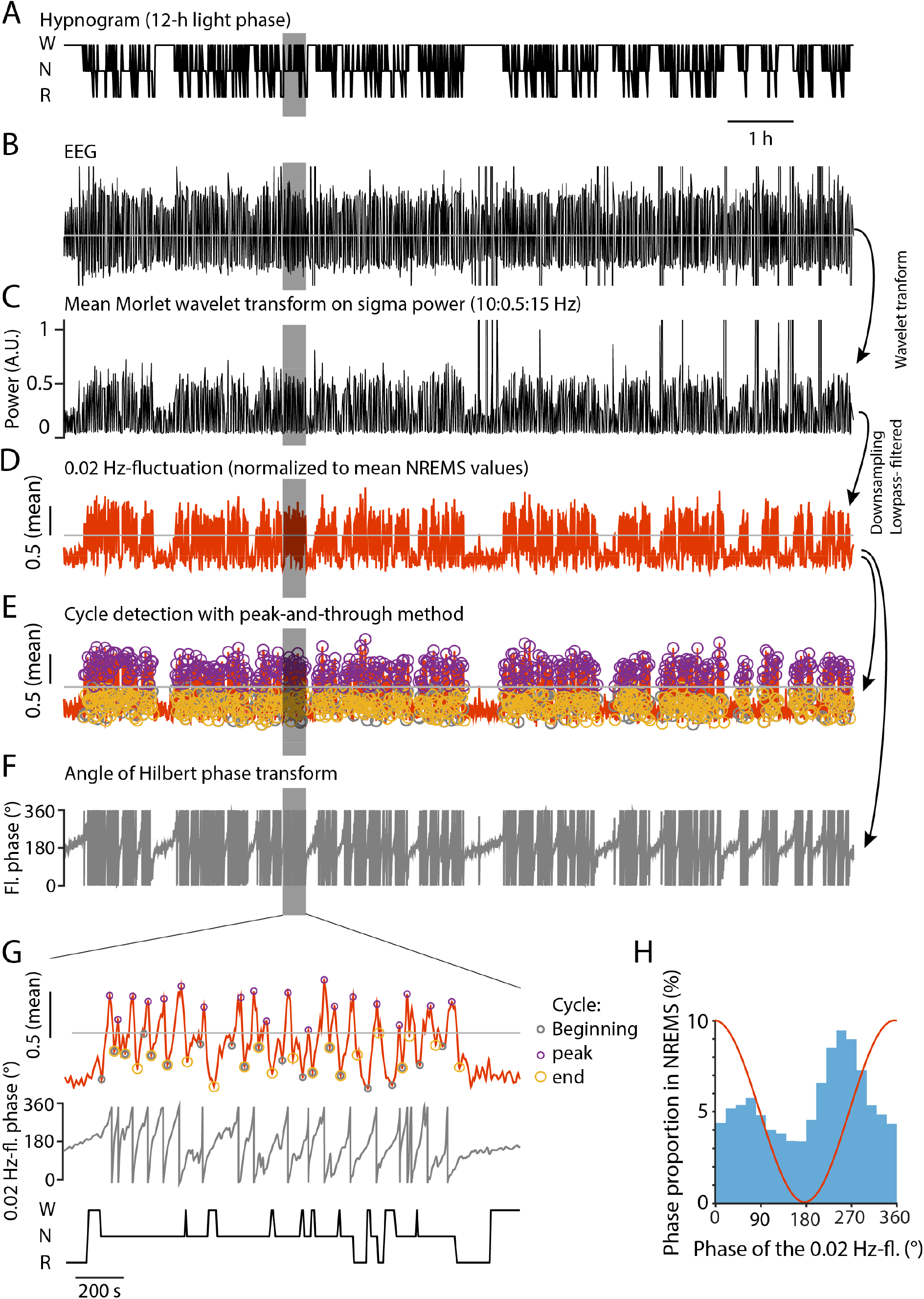
Methodological illustration of how to extract the 0.02 Hz-fluctuation from raw signals, to detect individual cycles, and to calculate the angles of the Hilbert transform. (**A**) 12-h hypnogram obtained from visual scoring of EEG/EMG recordings. W, wakefulness; N, NREMS; R, REMS. (**B**) Corresponding EEG signal. (**C**) Corresponding mean Morlet wavelet transform, calculated in 0.5-Hz frequency bins from 10—15 Hz. (**D**) Corresponding 0.02 Hz-fluctuation, obtained through downsampling to 10 Hz and lowpass filtering. Normalization was done by dividing the signal by its mean value in NREMS. **(E)** Result of cycle detection on the signal shown in D, using the peak-and-trough detection approach described in the Methods. The beginning, peak and end of each cycle is shown with color-coded circles. **(F)** Angle of the Hilbert transform of the signal shown D. The values are wrapped around 360° with 180° representing the troughs of the fluctuation. (**G**) Expansion of the grey area highlighted for D, E, F and A.(**H**) Histogram showing the phase constitution of the 0.02 Hz-fluctuation. Note the prominence from 180— 360°, indicating more time spent in ascending period compared to descending. A sinusoid would yield the same amount of points within each phase bin. Therefore, we now talk about 0.02 Hz-fluctuation and not oscillation as in (Lecci et al., 2017).

**Figure 5 – figure supplement 2.**
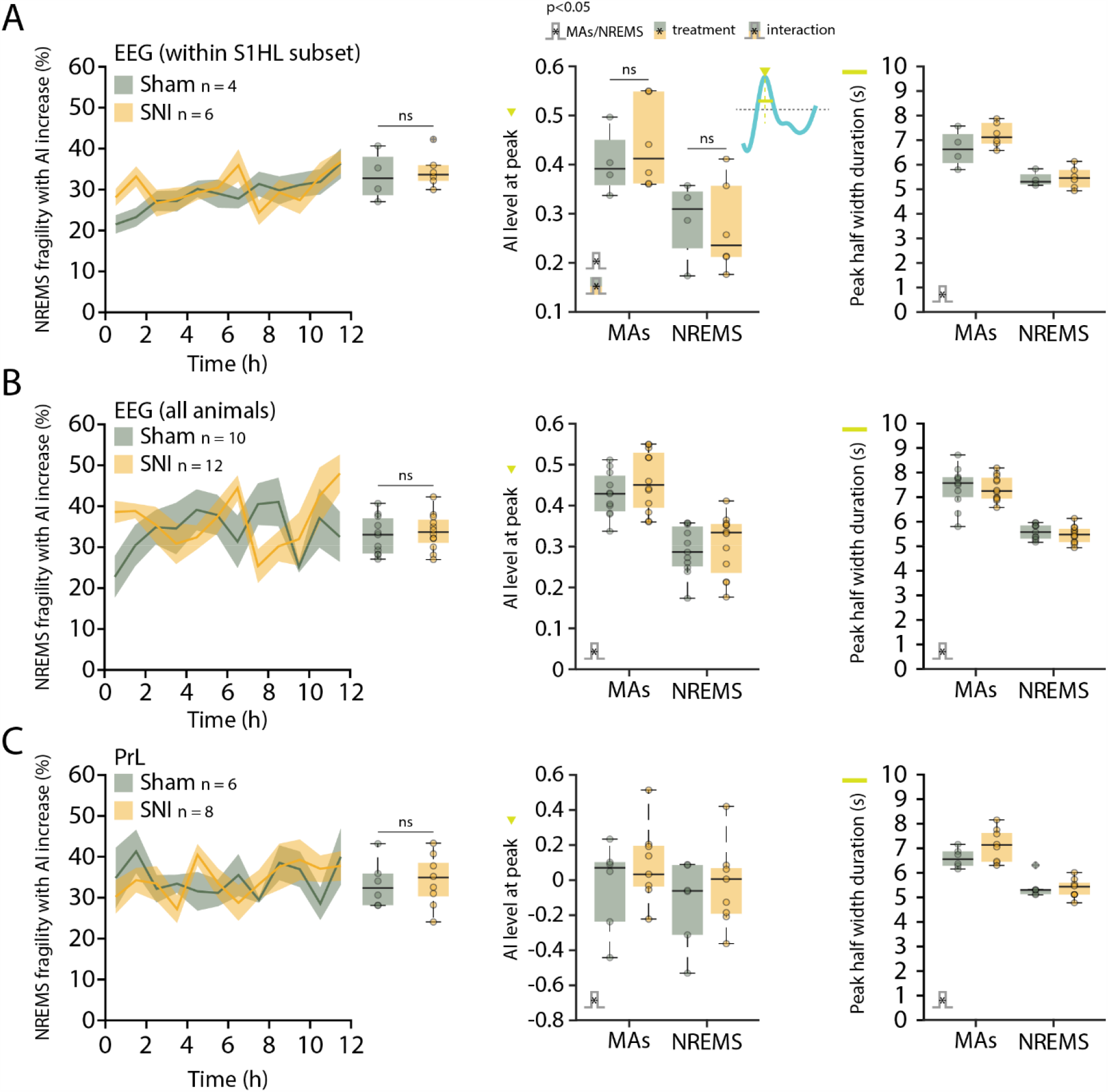
Activation index (AI) with peaks in fragility period are also present in EEG and PrL but occur in equal amounts in Sham or SNI. (**A**) Left, Proportion of fragility periods containing an AI peak over the 12-h light phase, detected in the frontoparietal EEG (contralateral to Sham or SNI surgeries) in the subset of animals from which the S1HL data were obtained in figure 5; sum rank test Sham vs SNI: W = 10, p = 0.76. The data from 1 Sham and 3 SNI animals were left out because the EEG was implanted ipsilateral to Sham or SNI surgeries. Middle: mean amplitude (green triangle in the inset) of the AI peaks in the EEG (S1HL subset), with or without the presence of a MA in the fragility period, in Sham and SNI. Mixed-model ANOVA: F_(1,8)_ = 0.03, p = 0.85 for ‘treatment’; F_(1,8)_ = 224.9, p = 3.8×10^−7^ for ‘MA’; F_(1,8)_ = 6.83, p = 0.03 for interaction. *Post-hoc* rank sum test for Sham *vs* SNI for fragility periods with a MA: W = 9, p = 0.6; for fragility periods without a MA: W = 13, p = 0.91; signed rank test with *vs* without MA in Sham: V = 10, p = 0.12; in SNI: V = 21, p = 0.03; α = 0.0125. Right: half width duration (green line in the inset) of the AI peaks in the EEG (S1HL subset), with or without the presence of a MA in the fragility period, in Sham and SNI. Mixed-model ANOVA: F_(1,8)_ = 2.06, p = 0.18 for ‘treatment’; F_(1,8)_ = 39.12, p = 2.4×10^−4^ for ‘MA’; F_(1,8)_ = 0.91, p = 0.36 for interaction. (**B**) Same layout as in A. Data from the frontoparietal EEG (contralateral to Sham or SNI surgeries) from all our animals; Left: proportion of fragility periods containing an AI peak, over time and 12 h quantification: unpaired *t*-tests Sham vs SNI: t_(21)_ = −0.5, p = 0.62. Middle: mean amplitude of the AI peaks in the EEG with or without the presence of a MA in the fragility period, in Sham and SNI. Mixed-model ANOVA: F_(1,21)_ = 0.81, p = 0.37 for ‘treatment’; F_(1,21)_ = 446.8, p = 1.23×10^−15^ for ‘MA’; F_(1,21)_ = 1.64, p = 0.21 for interaction. Right: duration at half-maximal amplitude of the AI peaks in the EEG, with or without the presence of a MA in the fragility period, in Sham and SNI. Mixed-model ANOVA: F_(1,21)_ = 0.29, p = 0.59 for ‘treatment’; F_(1,21)_ = 174.9, p = 1.2×10^−11^ for ‘MA’; F_(1,21)_ = 0.03, p = 0.85 for interaction. (**C**) Same layout as in A and B. Data from the PrL cortex; Left: proportion of fragility periods containing an AI peak, over time and 12 h quantification: unpaired *t*-tests Sham vs SNI: t_(12)_ = − 0.23, p = 0.82. Middle: mean amplitude of the AI peaks in PrL with or without the presence of a MA in the fragility period, in Sham and SNI. Mixed-model ANOVA: F_(1,12)_ = 0.89, p = 0.36 for ‘treatment’; F_(1,12)_ = 63.9, p = 3.7×10^−6^ for ‘MA’; F_(1,12)_ = 0.74, p = 0.4 for interaction. Right: duration at half-maximal amplitude of the AI peaks in the EEG, with or without the presence of a MA in the fragility period, in Sham and SNI. Mixed-model ANOVA: F_(1,12)_ = 1.84, p = 0.19 for ‘treatment’; F_(1,12)_ = 50.26, p = 1.2×10^−5^ for ‘MA’; F_(1,12)_ = 1.71, p = 0.21 for interaction.

**Figure 6 – figure supplement 3.**
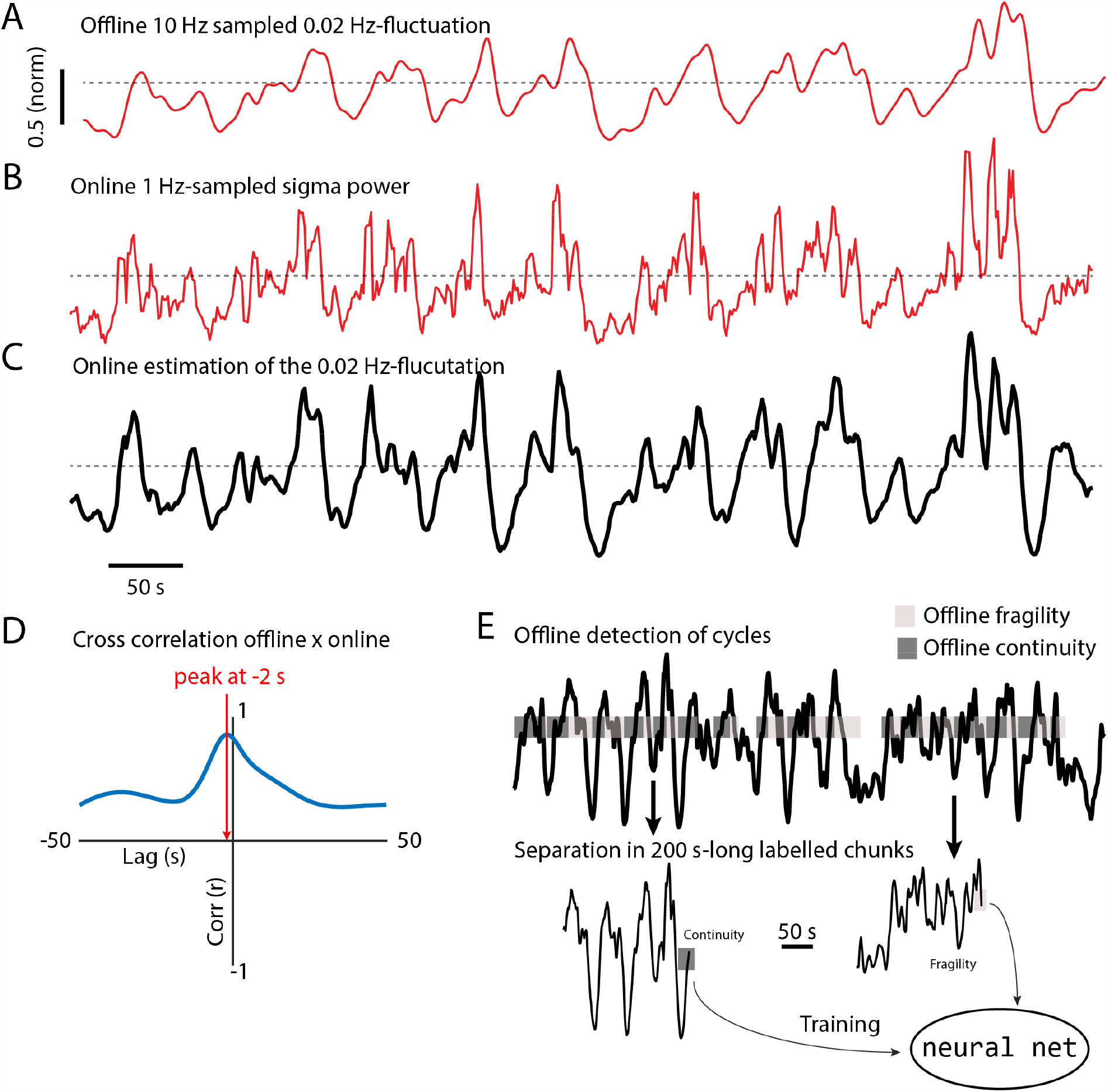
Online 0.02 Hz-fluctuation extraction and training of the neural network. (**A**) 0.02 Hz-fluctuation from a NREMS period extracted offline as described in ***Figure 2 − figure supplement 1***. (**B**) Corresponding sigma power values collected online at 1 Hz. (**C**) Corresponding online-estimated 0.02 Hz-fluctuation, based on a 9^th^ order polynomial fit on the sigma values shown in B. (**D**) Validation of the online estimate through cross correlation with offline data from a 12 h recording from one animal. The red arrow indicates the position of the highest correlation, showing a minor lag (−2 s). (**E**) Illustration of data labeling to train the neural network. The cycle detection (see method) was used offline on the 0.02 Hz-fluctuation previously extracted online. Continuity was set to the ascending periods and fragility to the descending one. When conditions for complete cycles were not met, the label ‘none’ was applied. Then, 200 s-long chunks of this signal were extracted and labelled with continuity, fragility or none depending on the position of the last point.

**Figure 7 – figure supplement 4.**
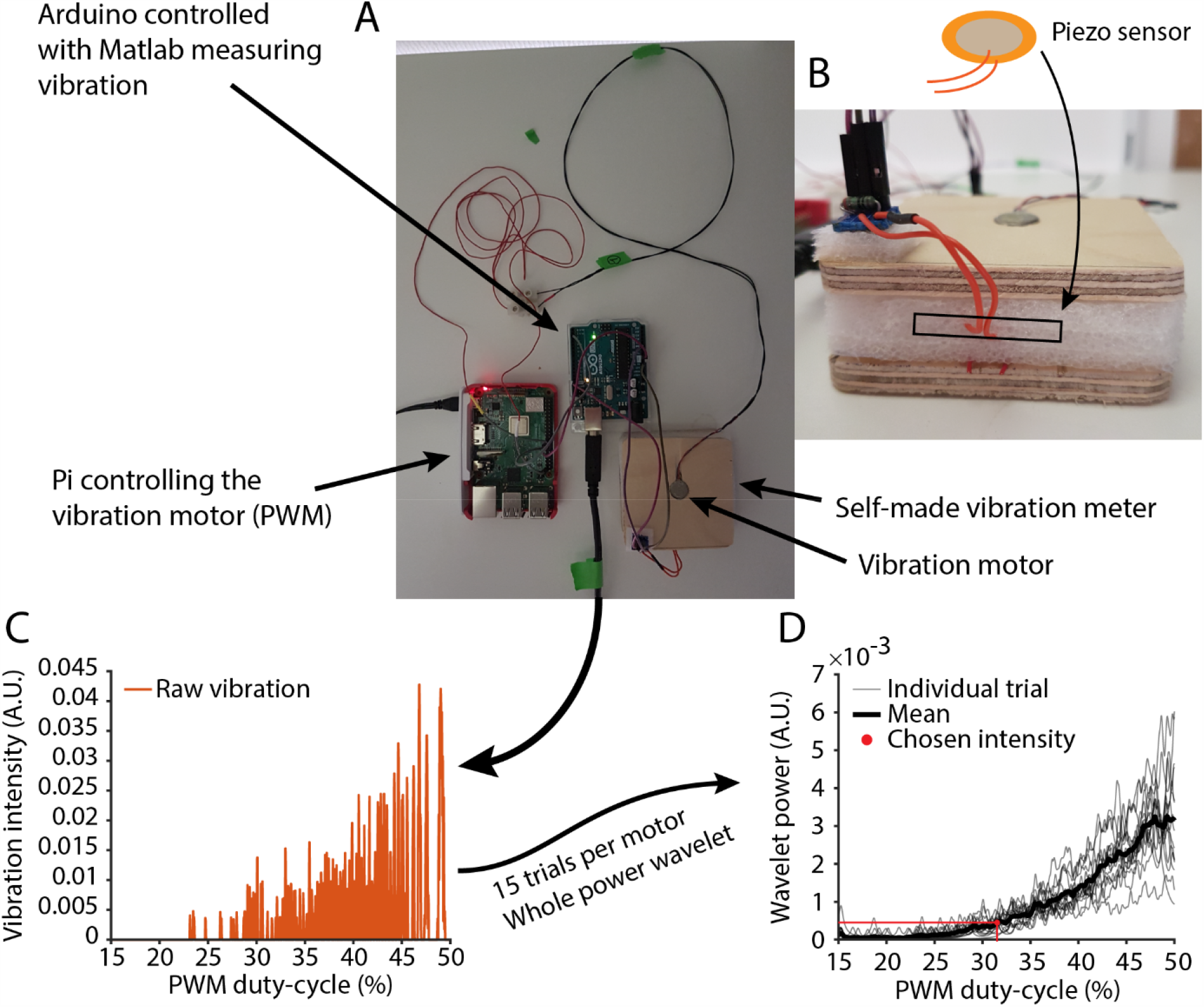
Vibration motor calibration for closed-loop somatosensory arousability testing. (**A**) Photo overview of the calibration setup, showing the vibration motor positioned onto a self-made piezometer containing the sensor (bottom right). The analog voltage generated by the sensor was measured by an Arduino and data fed into Matlab. To start vibrations, a raspberry pi sent a PWM signal to the vibration motor to modulate vibration intensity (achieved through increasing duty cycle from 15 to 50 %). A TTL signal sent from the raspberry pi to the Arduino initiated the trial. (**B**) Close-up view of the self-made piezometer. The piezometer was composed of protective foam sandwiched and glued between two thin wooden plates. The upper plate contained a hole the size of the vibration motors and the lower one was fixed to the table using two-sided adhesive tape. The piezo sensor was inserted inside the foam part and a resistance (1000 Ohm) was used in the circuit. (**C**) Plot of data received from one vibration measurement trial. The vibration curve was calculated from each trial using whole-power wavelet transform. (**D**) Calibration curve (mean of 15 trials) used to find the necessary duty-cycle value. The chosen intensity was the same for all the motors.

## References

Alexandre, C., Latremoliere, A., Ferreira, A., Miracca, G., Yamamoto, M., Scammell, T. E., & Woolf, C. J. (2017). Decreased alertness due to sleep loss increases pain sensitivity in mice. Nat Med, 23(6), 768–774. doi: 10.1038/nm.4329

Andersen, M. L., & Tufik, S. (2003). Sleep patterns over 21-day period in rats with chronic constriction of sciatic nerve. Brain Res, 984(1-2), 84–92. doi: 10.1016/s0006-8993(03)03095-6

Aston-Jones, G., & Bloom, F. E. (1981). Norepinephrine-containing locus coeruleus neurons in behaving rats exhibit pronounced responses to non-noxious environmental stimuli. J Neurosci, 1(8), 887–900.

Azarbarzin, A., Ostrowski, M., Hanly, P., & Younes, M. (2014). Relationship between arousal intensity and heart rate response to arousal. Sleep, 37(4), 645–653. doi: 10.5665/sleep.3560

Berens, P. (2009). CircStat: A MATLAB toolbox for circular statistics. J Stat Software, 31(10), 1–21. doi: 10.18637/jss.v031.i10

Bergmann, B. M., Winter, J. B., Rosenberg, R. S., & Rechtschaffen, A. (1987). NREM sleep with low-voltage EEG in the rat. Sleep, 10(1), 1–11.

Bjurstrom, M. F., & Irwin, M. R. (2016). Polysomnographic characteristics in nonmalignant chronic pain populations: A review of controlled studies. Sleep Med Rev, 26, 74–86. doi: 10.1016/j.smrv.2015.03.004

Bonnet, M., Carley, D., Carskadon, M., Easton, P., Guilleminault, C., Harper, R., Hayes, B., Hirshkowitz, M., Ktonas, P., Keenan, S., Pressman, M., Roehrs, T., Smith, J., Walsh, J., Weber, S., & Westbrook, P. (1992). The Atlas Task Force. EEG arousals: Scoring rules and examples. Sleep, 15, 173–184.

Bourquin, A. F., Süveges, M., Pertin, M., Gilliard, N., Sardy, S., Davison, A. C., Spahn, D. R., & Décosterd, I. (2006). Assessment and analysis of mechanical allodynia-like behavior induced by spared nerve injury (SNI) in the mouse. Pain, 122(1-2), 14 e11–14. doi: 10.1016/j.pain.2005.10.036

Burma, N. E., Leduc-Pessah, H., Fan, C. Y., & Trang, T. (2017). Animal models of chronic pain: Advances and challenges for clinical translation. J Neurosci Res, 95(6), 1242–1256. doi: 10.1002/jnr.23768

Buysse, D. J., Germain, A., Hall, M. L., Moul, D. E., Nofzinger, E. A., Begley, A., Ehlers, C. L., Thompson, W., & Kupfer, D. J. (2008). EEG spectral analysis in primary insomnia: NREM period effects and sex differences. Sleep, 31(12), 1673–1682. doi: 10.1093/sleep/31.12.1673

Cano, G., Mochizuki, T., & Saper, C. B. (2008). Neural circuitry of stress-induced insomnia in rats. J Neurosci, 28(40), 10167–10184. doi: 10.1523/JNEUROSCI.1809-08.2008

Cardoso-Cruz, H., Sameshima, K., Lima, D., & Galhardo, V. (2011). Dynamics of circadian thalamocortical flow of information during a peripheral neuropathic pain condition. Front Integr Neurosci, 5, 43. doi: 10.3389/fnint.2011.00043

Christensen, J. A. E., Wassing, R., Wei, Y., Ramautar, J. R., Lakbila-Kamal, O., Jennum, P. J., & Van Someren, E. J. W. (2019). Data-driven analysis of EEG reveals concomitant superficial sleep during deep sleep in insomnia disorder. Front Neurosci, 13, 598. doi: 10.3389/fnins.2019.00598

Claude, L., Chouchou, F., Prados, G., Castro, M., De Blay, B., Perchet, C., Garcia-Larrea, L., Mazza, S., & Bastuji, H. (2015). Sleep spindles and human cortical nociception: a surface and intracerebral electrophysiological study. J Physiol, 593(22), 4995–5008. doi: 10.1113/JP270941

Decosterd, I., & Woolf, C. J. (2000). Spared nerve injury: an animal model of persistent peripheral neuropathic pain. Pain, 87(2), 149–158. doi: 10.1016/s0304-3959(00)00276-1

Devor, M. (2009). Ectopic discharge in Aβ afferents as a source of neuropathic pain. Exp Brain Res, 196(1), 115–128. doi: 10.1007/s00221-009-1724-6

Dvir, H., Elbaz, I., Havlin, S., Appelbaum, L., Ivanov, P. C., & Bartsch, R. P. (2018). Neuronal noise as an origin of sleep arousals and its role in sudden infant death syndrome. Sci Adv, 4(4), eaar6277. doi: 10.1126/sciadv.aar6277

Feige, B., Baglioni, C., Spiegelhalder, K., Hirscher, V., Nissen, C., & Riemann, D. (2013). The microstructure of sleep in primary insomnia: an overview and extension. Int J Psychophysiol, 89(2), 171–180. doi: 10.1016/j.ijpsycho.2013.04.002

Feige, B., Nanovska, S., Baglioni, C., Bier, B., Cabrera, L., Diemers, S., Quellmalz, M., Siegel, M., Xeni, I., Szentkiralyi, A., Doerr, J. P., & Riemann, D. (2018). Insomnia-perchance a dream? Results from a NREM/REM sleep awakening study in good sleepers and patients with insomnia. Sleep, 41(5). doi: 10.1093/sleep/zsy032

Fernandez, L. M., Vantomme, G., Osorio-Forero, A., Cardis, R., Béard, E., & Lüthi, A. (2018). Thalamic reticular control of local sleep in sensory cortex. Elife, e39111. doi: doi: 10.7554/eLife.39111

Fernandez, L. M. J., Lecci, S., Cardis, R., Vantomme, G., Béard, E., & Lüthi, A. (2017). Quantifying infra-slow dynamics of spectral power and heart rate in sleeping mice. J Vis Exp(126). doi: 10.3791/55863

Fernandez, L. M. J., & Lüthi, A. (2020). Sleep Spindles: Mechanisms and Functions. Physiol Rev, 100(2), 805–868. doi: 10.1152/physrev.00042.2018

Finnerup, N. B., Attal, N., Haroutounian, S., McNicol, E., Baron, R., Dworkin, R. H., Gilron, I., Haanpaa, M., Hansson, P., Jensen, T. S., Kamerman, P. R., Lund, K., Moore, A., Raja, S. N., Rice, A. S., Rowbotham, M., Sena, E., Siddall, P., Smith, B. H., & Wallace, M. (2015). Pharmacotherapy for neuropathic pain in adults: a systematic review and meta-analysis. Lancet Neurol, 14(2), 162–173. doi: 10.1016/S1474-4422(14)70251-0

Finnerup, N. B., Kuner, R., & Jensen, T. S. (2021). Neuropathic pain: from mechanisms to treatment. Physiol Rev, 101(1), 259–301. doi: 10.1152/physrev.00045.2019

Forget, D., Morin, C. M., & Bastien, C. H. (2011). The role of the spontaneous and evoked K-complex in good-sleeper controls and in individuals with insomnia. Sleep, 34(9), 1251–1260. doi: 10.5665/SLEEP.1250

Franken, P. (2002). Long-term vs. short-term processes regulating REM sleep. J Sleep Res, 11(1), 17–28. Franken, P., Malafosse, A., & Tafti, M. (1999). Genetic determinants of sleep regulation in inbred mice. Sleep, 22(2), 155–169.

Fulda, S. (2011). Idiopathic REM sleep behavior disorder as a long-term predictor of neurodegenerative disorders. EPMA J, 2(4), 451–458. doi: 10.1007/s13167-011-0096-8

Guida, F., De Gregorio, D., Palazzo, E., Ricciardi, F., Boccella, S., Belardo, C., Iannotta, M., Infantino, R., Formato, F., Marabese, I., Luongo, L., de Novellis, V., & Maione, S. (2020). Behavioral, biochemical and electrophysiological changes in spared nerve injury model of neuropathic pain. Int J Mol Sci, 21(9). doi: 10.3390/ijms21093396

Hayat, H., Regev, N., Matosevich, N., Sales, A., Paredes-Rodriguez, E., Krom, A. J., Bergman, L., Li, Y., Lavigne, M., Kremer, E. J., Yizhar, O., Pickering, A. E., & Nir, Y. (2020). Locus coeruleus norepinephrine activity mediates sensory-evoked awakenings from sleep. Sci Adv, 6(15), eaaz4232. doi: 10.1126/sciadv.aaz4232

Hohn, V. D., May, E. S., & Ploner, M. (2019). From correlation towards causality: modulating brain rhythms of pain using transcranial alternating current stimulation. Pain Rep, 4(4), e723. doi: 10.1097/PR9.0000000000000723

Huang, Z. L., Mochizuki, T., Qu, W. M., Hong, Z. Y., Watanabe, T., Urade, Y., & Hayaishi, O. (2006). Altered sleep-wake characteristics and lack of arousal response to H3 receptor antagonist in histamine H1 receptor knockout mice. Proc Natl Acad Sci U S A, 103(12), 4687–4692. doi: 10.1073/pnas.0600451103

Iber, C., Ancoli-Israel, S., Chesson, A., & Quan, S. F. (2007). The AASM manual for the scoring of sleep and associated events: rules, terminology and technical specifications. Westchester, IL: American Academy of Sleep Medicine.

Kjaerby, C., Andersen, M., Hauglund, N., Ding, F., Wang, W., Xu, Q., Deng, S., Kang, N., Peng, S., Sun, Q., Dall, C., Jørgensen, P. K., Feng, J., Li, Y., Weikop, P., Hirase, H., & Nedergaard, M. (2020). Dynamic fluctuations of the locus coeruleus-norepinephrine system underlie sleep state transitions. bioRxiv, 2020.2009.2001.274977. doi: 10.1101/2020.09.01.274977

Kontinen, V. K., Ahnaou, A., Drinkenburg, W. H., & Meert, T. F. (2003). Sleep and EEG patterns in the chronic constriction injury model of neuropathic pain. Physiol Behav, 78(2), 241–246. doi: 10.1016/s0031-9384(02)00966-6

Kopp, C., Longordo, F., Nicholson, J. R., & Lüthi, A. (2006). Insufficient sleep reversibly alters bidirectional synaptic plasticity and NMDA receptor function. J Neurosci, 26(48), 12456–12465. doi: 10.1523/JNEUROSCI.2702-06.2006

Krystal, A. D., & Edinger, J. D. (2008). Measuring sleep quality. Sleep Med, 9 Suppl 1, S10–17. doi: 10.1016/S1389-9457(08)70011-X

Krystal, A. D., Edinger, J. D., Wohlgemuth, W. K., & Marsh, G. R. (2002). NREM sleep EEG frequency spectral correlates of sleep complaints in primary insomnia subtypes. Sleep, 25(6), 630–640.

Kuner, R., & Kuner, T. (2020). Cellular circuits in the brain and their modulation in acute and chronic pain. Physiol Rev. doi: 10.1152/physrev.00040.2019

Landis, C. A., Levine, J. D., & Robinson, C. R. (1989). Decreased slow-wave and paradoxical sleep in a rat chronic pain model. Sleep, 12(2), 167–177. doi: 10.1093/sleep/12.2.167

LeBlanc, B. W., Lii, T. R., Silverman, A. E., Alleyne, R. T., & Saab, C. Y. (2014). Cortical theta is increased while thalamocortical coherence is decreased in rat models of acute and chronic pain. Pain, 155(4), 773–782. doi: 10.1016/j.pain.2014.01.013

Lecci, S., Cataldi, J., Betta, M., Bernardi, G., Heinzer, R., & Siclari, F. (2020). EEG changes associated with subjective under- and overestimation of sleep duration. Sleep, 43, zsaa094. doi: 10.1093/sleep/zsaa094

Lecci, S., Fernandez, L. M., Weber, F. D., Cardis, R., Chatton, J. Y., Born, J., & Lüthi, A. (2017). Coordinated infraslow neural and cardiac oscillations mark fragility and offline periods in mammalian sleep. Sci Adv, 3(2), e1602026. doi: 10.1126/sciadv.1602026

Léna, C., Popa, D., Grailhe, R., Escourrou, P., Changeux, J. P., & Adrien, J. (2004). β2-containing nicotinic receptors contribute to the organization of sleep and regulate putative micro-arousals in mice. J Neurosci, 24(25), 5711–5718. doi: 10.1523/JNEUROSCI.3882-03.2004

Leys, L. J., Chu, K. L., Xu, J., Pai, M., Yang, H. S., Robb, H. M., Jarvis, M. F., Radek, R. J., & McGaraughty, S. (2013). Disturbances in slow-wave sleep are induced by models of bilateral inflammation, neuropathic, and postoperative pain, but not osteoarthritic pain in rats. Pain, 154(7), 1092–1102. doi: 10.1016/j.pain.2013.03.019

Li, S. B., Borniger, J. C., Yamaguchi, H., Hedou, J., Gaudilliere, B., & de Lecea, L. (2020). Hypothalamic circuitry underlying stress-induced insomnia and peripheral immunosuppression. Sci Adv, 6(37). doi: 10.1126/sciadv.abc2590

Lo, C. C., Chou, T., Penzel, T., Scammell, T. E., Strecker, R. E., Stanley, H. E., & Ivanov, P. (2004). Common scale-invariant patterns of sleep-wake transitions across mammalian species. Proc Natl Acad Sci U S A, 101(50), 17545–17548. doi: 10.1073/pnas.0408242101

Maes, J., Verbraecken, J., Willemen, M., De Volder, I., van Gastel, A., Michiels, N., Verbeek, I., Vandekerckhove, M., Wuyts, J., Haex, B., Willemen, T., Exadaktylos, V., Bulckaert, A., & Cluydts, R. (2014). Sleep misperception, EEG characteristics and autonomic nervous system activity in primary insomnia: a retrospective study on polysomnographic data. Int J Psychophysiol, 91(3), 163–171. doi: 10.1016/j.ijpsycho.2013.10.012

Mang, G. M., & Franken, P. (2012). Sleep and EEG phenotyping in mice. Curr Protoc Mouse Biol, 2(1), 55–74. doi: 10.1002/9780470942390.mo110126

Mathias, J. L., Cant, M. L., & Burke, A. L. J. (2018). Sleep disturbances and sleep disorders in adults living with chronic pain: a meta-analysis. Sleep Med, 52, 198–210. doi: 10.1016/j.sleep.2018.05.023

May, E. S., Nickel, M. M., Ta Dinh, S., Tiemann, L., Heitmann, H., Voth, I., Tolle, T. R., Gross, J., & Ploner, M. (2019). Prefrontal gamma oscillations reflect ongoing pain intensity in chronic back pain patients. Hum Brain Mapp, 40(1), 293–305. doi: 10.1002/hbm.24373

Moisset, X., & Bouhassira, D. (2007). Brain imaging of neuropathic pain. Neuroimage, 37 Suppl 1, S80–88. doi: 10.1016/j.neuroimage.2007.03.054

Moldofsky, H., Scarisbrick, P., England, R., & Smythe, H. (1975). Musculoskeletal symptoms and non-REM sleep disturbance in patients with “fibrositis syndrome” and healthy subjects. Psychosom Med, 37, 341–351.

Neckelmann, D., & Ursin, R. (1993). Sleep stages and EEG power spectrum in relation to acoustical stimulus arousal threshold in the rat. Sleep, 16(5), 467–477.

Nickel, M. M., May, E. S., Tiemann, L., Postorino, M., Ta Dinh, S., & Ploner, M. (2017). Autonomic responses to tonic pain are more closely related to stimulus intensity than to pain intensity. Pain, 158(11), 2129–2136. doi: 10.1097/j.pain.0000000000001010

Nobili, L., Ferrara, M., Moroni, F., De Gennaro, L., Russo, G. L., Campus, C., Cardinale, F., & De Carli, F. (2011). Dissociated wake-like and sleep-like electro-cortical activity during sleep. Neuroimage, 58(2), 612–619. doi: 10.1016/j.neuroimage.2011.06.032

Nollet, M., Hicks, H., McCarthy, A. P., Wu, H., Moller-Levet, C. S., Laing, E. E., Malki, K., Lawless, N., Wafford, K. A., Dijk, D. J., & Winsky-Sommerer, R. (2019). REM sleep’s unique associations with corticosterone regulation, apoptotic pathways, and behavior in chronic stress in mice. Proc Natl Acad Sci U S A, 116(7), 2733–2742. doi: 10.1073/pnas.1816456116

Parrino, L., Milioli, G., De Paolis, F., Grassi, A., & Terzano, M. G. (2009). Paradoxical insomnia: the role of CAP and arousals in sleep misperception. Sleep Med, 10(10), 1139–1145. doi: 10.1016/j.sleep.2008.12.014

Perlis, M. L., Merica, H., Smith, M. T., & Giles, D. E. (2001a). Beta EEG activity and insomnia. Sleep Med Rev, 5(5), 363–374. doi: 10.1053/smrv.2001.0151

Perlis, M. T., Smith, M. T., Andrews, P. J., Orff, H., & Giles, D. E. (2001b). Beta/Gamma EEG activity in patients with primary and secondary insomnia and good sleeper controls. Sleep, 24(1), 110–117. doi: 10.1093/sleep/24.1.110

Ploner, M., Sorg, C., & Gross, J. (2017). Brain rhythms of pain. Trends Cogn Sci, 21(2), 100–110. doi: 10.1016/j.tics.2016.12.001

Radzicki, D., Pollema-Mays, S. L., Sanz-Clemente, A., & Martina, M. (2017). Loss of M1 receptor dependent cholinergic excitation contributes to mPFC deactivation in neuropathic pain. J Neurosci, 37(9), 2292–2304. doi: 10.1523/JNEUROSCI.1553-16.2017

Riedner, B. A., Goldstein, M. R., Plante, D. T., Rumble, M. E., Ferrarelli, F., Tononi, G., & Benca, R. M. (2016). Regional patterns of elevated alpha and high-frequency electroencephalographic activity during nonrapid eye movement sleep in chronic insomnia: a pilot study. Sleep, 39(4), 801–812. doi: 10.5665/sleep.5632

Rolls, A., Colas, D., Adamantidis, A., Carter, M., Lanre-Amos, T., Heller, H. C., & de Lecea, L. (2011). Optogenetic disruption of sleep continuity impairs memory consolidation. Proc Natl Acad Sci U S A, 108(32), 13305–13310. doi: 10.1073/pnas.1015633108

Salin-Pascual, R. J., Roehrs, T. A., Merlotti, L. A., Zorick, F., & Roth, T. (1992). Long-term study of the sleep of insomnia patients with sleep state misperception and other insomnia patients. Am J Psychiatry, 149(7), 904–908. doi: 10.1176/ajp.149.7.904

Sarnthein, J., Stern, J., Aufenberg, C., Rousson, V., & Jeanmonod, D. (2006). Increased EEG power and slowed dominant frequency in patients with neurogenic pain. Brain, 129(Pt 1), 55–64. doi: 10.1093/brain/awh631

Schulz, E., May, E. S., Postorino, M., Tiemann, L., Nickel, M. M., Witkovsky, V., Schmidt, P., Gross, J., & Ploner, M. (2015). Prefrontal gamma oscillations encode tonic pain in humans. Cereb Cortex, 25(11), 4407–4414. doi: 10.1093/cercor/bhv043

Sforza, E., Jouny, C., & Ibanez, V. (2000). Cardiac activation during arousal in humans: further evidence for hierarchy in the arousal response. Clin Neurophysiol, 111(9), 1611–1619. doi: 10.1016/s1388-2457(00)00363-1

Shirvalkar, P., Veuthey, T. L., Dawes, H. E., & Chang, E. F. (2018). Closed-loop deep brain stimulation for refractory chronic pain. Front Comput Neurosci, 12, 18. doi: 10.3389/fncom.2018.00018

Silva, A., Andersen, M. L., & Tufik, S. (2008). Sleep pattern in an experimental model of osteoarthritis. Pain, 140(3), 446–455. doi: 10.1016/j.pain.2008.09.025

Silvani, A. (2019). Sleep disorders, nocturnal blood pressure, and cardiovascular risk: A translational perspective. Auton Neurosci, 218, 31–42. doi: 10.1016/j.autneu.2019.02.006

Spiegelhalder, K., Regen, W., Feige, B., Holz, J., Piosczyk, H., Baglioni, C., Riemann, D., & Nissen, C. (2012). Increased EEG sigma and beta power during NREM sleep in primary insomnia. Biol Psychol, 91(3), 329–333. doi: 10.1016/j.biopsycho.2012.08.009

St-Jean, G., Turcotte, I., & Bastien, C. H. (2012). Cerebral asymmetry in insomnia sufferers. Front Neurol, 3, 47. doi: 10.3389/fneur.2012.00047

Tan, L. L., Oswald, M. J., Heinl, C., Retana Romero, O. A., Kaushalya, S. K., Monyer, H., & Kuner, R. (2019). Gamma oscillations in somatosensory cortex recruit prefrontal and descending serotonergic pathways in aversion and nociception. Nat Commun, 10(1), 983. doi: 10.1038/s41467-019-08873-z

Tokunaga, S., Takeda, Y., Shinomiya, K., Yamamoto, W., Utsu, Y., Toide, K., & Kamei, C. (2007). Changes of sleep patterns in rats with chronic constriction injury under aversive conditions. Biol Pharm Bull, 30(11), 2088–2090. doi: 10.1248/bpb.30.2088

Treede, R. D., Rief, W., Barke, A., Aziz, Q., Bennett, M. I., Benoliel, R., Cohen, M., Evers, S., Finnerup, N. B., First, M. B., Giamberardino, M. A., Kaasa, S., Korwisi, B., Kosek, E., Lavand’homme, P., Nicholas, M., Perrot, S., Scholz, J., Schug, S., Smith, B. H., Svensson, P., Vlaeyen, J. W. S., & Wang, S. J. (2019). Chronic pain as a symptom or a disease: the IASP Classification of Chronic Pain for the International Classification of Diseases (ICD-11). Pain, 160(1), 19–27. doi: 10.1097/j.pain.0000000000001384

van Someren, E. J. W. (2020). Brain mechanisms of insomnia: new perspectives on causes and consequences. Physiol Rev, in press.

Vargas, I., Nguyen, A. M., Muench, A., Bastien, C. H., Ellis, J. G., & Perlis, M. L. (2020). Acute and chronic insomnia: what has time and/or hyperarousal got to do with it? Brain Sci, 10(2). doi: 10.3390/brainsci10020071

Vassalli, A., & Franken, P. (2017). Hypocretin (orexin) is critical in sustaining theta/gamma-rich waking behaviors that drive sleep need. Proc Natl Acad Sci U S A, 114(27), E5464–E5473. doi: 10.1073/pnas.1700983114

Wall, P. D., & Devor, M. (1983). Sensory afferent impulses originate from dorsal root ganglia as well as from the periphery in normal and nerve injured rats. Pain, 17(4), 321–339. doi: 10.1016/0304-3959(83)90164-1

Wei, Y., Colombo, M. A., Ramautar, J. R., Blanken, T. F., van der Werf, Y. D., Spiegelhalder, K., Feige, B., Riemann, D., & Van Someren, E. J. W. (2017). Sleep stage transition dynamics reveal specific stage 2 vulnerability in insomnia. Sleep, 40(9). doi: 10.1093/sleep/zsx117

Wimmer, R. D., Astori, S., Bond, C. T., Rovó, Z., Chatton, J. Y., Adelman, J. P., Franken, P., & Lüthi, A. (2012). Sustaining sleep spindles through enhanced SK2-channel activity consolidates sleep and elevates arousal threshold. J Neurosci, 32(40), 13917–13928. doi: 10.1523/JNEUROSCI.2313-12.2012

Yüzgeç, O., Prsa, M., Zimmermann, R., & Huber, D. (2018). Pupil size coupling to cortical states protects the stability of deep sleep via parasympathetic modulation. Curr Biol, 28(3), 392–400 e393. doi: 10.1016/j.cub.2017.12.049

Zhang, Z., Gadotti, V. M., Chen, L., Souza, I. A., Stemkowski, P. L., & Zamponi, G. W. (2015). Role of prelimbic GABAergic circuits in sensory and emotional aspects of neuropathic pain. Cell Rep, 12(5), 752–759. doi: 10.1016/j.celrep.2015.07.001

